# Measuring genetic variation in the multi-ethnic Million Veteran Program (MVP)

**DOI:** 10.1101/2020.01.06.896613

**Authors:** Haley Hunter-Zinck, Yunling Shi, Man Li, Bryan R. Gorman, Sun-Gou Ji, Ning Sun, Teresa Webster, Andrew Liem, Paul Hsieh, Poornima Devineni, Purushotham Karnam, Lakshmi Radhakrishnan, Jeanette Schmidt, Themistocles L. Assimes, Jie Huang, Cuiping Pan, Donald Humphries, Mary Brophy, Jennifer Moser, Sumitra Muralidhar, Grant D. Huang, Ronald Przygodzki, John Concato, John M. Gaziano, Joel Gelernter, Christopher J. O’Donnell, Elizabeth R. Hauser, Hongyu Zhao, Timothy J. O’Leary, Philip S. Tsao, Saiju Pyarajan, on behalf of the VA Million Veteran Program

## Abstract

The Million Veteran Program (MVP), initiated by the Department of Veterans Affairs (VA), aims to collect consented biosamples from at least one million Veterans. Presently, blood samples have been collected from over 800,000 enrolled participants. The size and diversity of the MVP cohort, as well as the availability of extensive VA electronic health records make it a promising resource for precision medicine. MVP is conducting array-based genotyping to provide genome-wide scan of the entire cohort, in parallel with whole genome sequencing, methylation, and other omics assays. Here, we present the design and performance of MVP 1.0 custom Axiom^®^ array, which was designed and developed as a single assay to be used across the multi-ethnic MVP cohort. A unified genetic quality control analysis was developed and conducted on an initial tranche of 485,856 individuals leading to a high-quality dataset of 459,777 unique individuals. 668,418 genetic markers passed quality control and showed high quality genotypes not only on common variants but also on rare variants. We confirmed the substantial ancestral diversity of MVP with nearly 30% non-European individuals, surpassing other large biobanks. We also demonstrated the quality of the MVP dataset by replicating established genetic associations with height in European Americans and African Americans ancestries. This current data set has been made available to approved MVP researchers for genome-wide association studies and other downstream analyses. Further data releases will be available for analysis as recruitment at the VA continues and the cohort expands both in size and diversity.

## Introduction

The Department of Veterans Affairs (VA) initiated the Million Veteran Program (MVP) in 2011 to create a mega-biobank of at least one million samples with genetic data linked to nationally consolidated longitudinal clinical records^1^. The initial and continuing goal of MVP is to create a national resource for research to improve the health of United States Veterans and, more generally, to contribute to our understanding of human health. MVP has currently collected samples from over 800,000 Veteran participants and with continued recruitment efforts expects to exceed a total of 1 million participants in the next 2 to 3 years.

While MVP is similar in some respects to other large biobank projects such as the UK Biobank, the Kaiser Permanente Research Program on Genes, Environment, and Health (RPGEH), the China Kadoorie Biobank (CKB), and the DiscovEHR initiative^2–4^, it is unique in several ways. MVP is one of the largest single biobanking efforts to date, satisfying the need for larger genetic datasets while also benefiting from a very rich, nationally integrated longitudinal clinical database housed in the largest consolidated healthcare network in the United States. This feature allows for enhanced clinical phenotyping capabilities. The availability of additional self-reported health and lifestyle survey information augments clinical data from the Veterans Information Systems and Technology Architecture (VistA) – the VA’s Electronic Health Record (EHR).

Furthermore, with over 29% of participants self-reporting non-white ethnicity, MVP has substantial diversity in genetic ancestry, meeting a pressing need for greater diversity in genome-wide association analyses to discover novel associations, reduce false positives, and increase research equity^5–8^. As such, the MVP cohort provides an unprecedented opportunity for increasing the power of genome-wide association studies (GWAS) and will enable association discoveries for clinically important low frequency and rare variants possible only in larger sample sizes. Reliable typing of these variants may provide explanations of missing genetic susceptibility in complex or non-Mendelian diseases. However, the genetic diversity of MVP also poses challenges in genotype quality control.

In this report, we introduce the first installment of MVP genotype data consisting of 459,777 samples surveyed at 668,418 markers. In brief, we 1) describe the design of a research genotyping array with emphasis on clinically useful and/or rare variants applicable to multi-ethnic backgrounds; 2) describe the generation and quality control of genotyping data; 3) highlight some of the unique features of the current MVP dataset, including exploratory analyses of genetic ancestry; and 4) replicate effect sizes of previously reported variants associated with height in European Americans and African Americans. Overall, we find that the MVP genetic dataset, linked to deep phenotypic data, is a high-quality and diverse resource for performing genetic analyses.

## Materials and Methods

### Human subjects and data and sample collection

The VA Central Institutional Review Board (IRB), as well as the local IRBs at the VA Boston Healthcare System and the VA Connecticut Healthcare System, approved this project. An overview of the recruitment strategies and protocols is given in a previous publication^1^. Briefly, participants were recruited from approximately 60 VA healthcare facilities across the United States on a rolling basis. Informed consent was obtained from all participants. Participants consented to a blood draw and to have their DNA analyzed, as well as to linking their genetic information with their full clinical, survey and other health data. Participants were also invited to answer two separate surveys about basic demographic information and lifestyle characteristics.

Blood drawn from consenting participants was shipped to the central biorepository in Boston, Massachusetts where DNA was extracted and later shipped to two external vendors for genotyping on a custom Axiom^®^ array designed specifically for MVP (MVP 1.0). A description of the MVP 1.0 array design features is detailed in Supplementary Materials.

### Thermo Fisher Scientific (formally Affymetrix) Axiom^®^ Genotyping Platform

The MVP 1.0 custom Axiom^®^ array is based on the Axiom^®^ Genotyping Platform. The Axiom genotyping platform utilizes a two-color, ligation-based assay using 30-mer oligonucleotide probes synthesized *in situ* onto a microarray substrate. Each single nucleotide polymorphism (SNP) feature contains a unique oligomeric sequence complementary to the genomic sequence flanking the polymorphic site on either the forward or the reverse strand. Solution probes bearing attachment sites for one of two dyes depending on the 3’ (SNP-site) base (A or T, versus C or G) are hybridized to the target complex, followed by ligation for specificity. Oligonucleotide sequences complementary to the forward or reverse strands are referred to as probesets. A marker (SNP or indel) can be interrogated by the forward and/or reverse strand probeset.

For additional details of the Axiom^®^ Genotyping Platform, see the Supplemental Materials and Methods.

### Genotype calling

We received unprocessed Axiom^®^ genotype data for 485,856 unique samples assayed by two vendors, referred to as Vendor 1 and Vendor 2, and performed genotype calling in batches grouped by vendor and sample processing date. Using data provided by the vendors and generated from our internal genotype calling process (see Supplemental Materials and Methods for details), we first analyzed the standard Axiom^®^ genotype quality metrics and compared these metrics between the two vendors.

After calling genotypes, we applied an advanced normalization procedure for mitigating plate-to-plate variation developed in collaboration with ThermoFisher Scientific Inc. The procedure was applied selectively on a per-batch basis to probesets exhibiting high plate-to-plate variance. After plate normalization, we applied standard marker quality control procedures to clean and harmonize genotype calls across all the batches (Supplemental Materials and Methods), followed by advanced sample QC.

### Advanced sample QC

#### Sample contamination

To detect and mitigate sample contamination, we assessed heterozygosity with PLINK, version 1.9, by calculating the F coefficient and quarantining samples with an F coefficient of less than −0.1. We assessed excess relatedness by using the relatedness inference software KING, version 2.0, and quarantined samples having a kinship coefficient of at least 0.1 with 7 or more other samples within MVP. These samples had high dish QC (DQC) and low call rates and were outliers compared to the majority of samples in the MVP dataset (Figure S5D). Because a call rate below 98.5% correlated with excess sample heterozygosity or relatedness, we removed samples (15,436, or 3.00%) with call rates below this threshold^9^. All samples that were removed or quarantined from the current release of MVP data will be re-genotyped and included in the future data releases.

#### Sample mislabeling

We identified samples and plates demonstrating potential mislabeling issues by analyzing genotype concordance between intentional duplicate samples that were sent blinded to the vendors as new samples for genotyping. Of the 25,867 intentional duplicate pairs, only 211 (0.82%) pairs were highly discordant (greater than 1% discordance). Samples on plates with discordant intentional duplicate pairs were quarantined for further analysis and re-genotyping. We also removed both samples and plates if the duplicate pair had a relatedness coefficient of less than 0.45. These precautions were taken due to the concern of potential plate swaps and led to 9,975 samples being quarantined.

#### Sample misidentification

To discriminate between misidentified intentional duplicates (same samples intentionally genotyped twice), technical duplicates (controls repeatedly genotyped by vendors), and monozygotic twins, we calculated sample relatedness with the KING software, version 2.1^10^. Monozygotic twins were confirmed by cross-referencing EHR data. Pairs with birth dates differing by no more than one day and having unique participant identifiers and first names were considered verified monozygotic twin pairs. Unverified samples were quarantined as potentially mislabeled and will be re-genotyped.

#### Sex check

To confirm sample gender, we extracted markers genotyped on the X chromosome while excluding the pseudoautosomal region, used the sex-check command from PLINK, and compared the expected F coefficient on the X chromosome to the gender recorded in the sample’s EHR for all samples^11^. Participants whose reported gender differed from that inferred by PLINK were quarantined from subsequent analysis. We also removed remaining samples on plates with 4 or more gender mismatches to account for potential plate swaps. The threshold is relatively low because of the low percentage of females in our dataset.

### Advanced marker QC

#### Advanced marker QC pipeline

We implemented three main approaches to create the advanced marker QC pipeline: (1) exclude probeset calls from all batches for probesets that failed advanced QC tests; (2) exclude probeset calls in a given batch for which the probeset is not recommended; and (3) choose the best probeset per marker for markers interrogated by multiple probesets, and exclude probesets calls from all batches for the “not-best” probesets. Details of each steps of the advanced marker QC are available in Supplemental Materials and Methods and in Figure S4, Figure S6A, and Figure S7A.

The advanced marker QC pipeline produced an inclusion list of probesets that met quality standards across the entire MVP dataset. For each batch, we included a probeset in the dataset if it met all three criteria: 1) included in the inclusion list; 2) recommended in that batch; and 3) was the best probeset for a marker interrogated by multiple probesets. We then generated a list of probesets per batch, created PLINK marker list binary files for each batch, and then merged all batches together using the PLINK merge command.

#### Reproducibility of genotype calling

To assess the consistency of genotype calls across time and vendors, we analyzed the discordance between 25,867 intentional duplicate samples that were sent to the vendors blinded. After confirming these sample pairs were genetically identical through KING relatedness inference, we determined the number of minor allele pairs (MAPs) for each marker. A MAP is any pair of genotypes for a marker where both pairs are called and the pair contains at least one minor allele. We then calculated the number of discordant genotyping pairs per MAP for each marker. Normalizing by the number of MAPs renders different MAF bins comparable in the discordance calculation. Otherwise, rare markers will always have extremely low discordance rates, as most samples carry the homozygous major genotype.

Additionally, within the 485,856 samples genotyped in the MVP cohort, we included 2,064 positive control samples. We called the genotypes of the positive controls along with other MVP samples across 112 batches organized by genotyping scan date for 668,418 markers passing advanced marker quality control. These genotypes were compared to the consensus positive control genotype.

To construct the consensus genotype sequence, we calculated the frequency of each marker across the panel of 2,064 positive control samples. Markers with MAF of less than 1% were set to homozygous in the consensus sequence, and markers with MAF of greater than 49% were set to heterozygous in the consensus sequence. For markers with MAF greater than or equal to 1% and less than or equal to 49% (536, or 0.082% of markers) or that had no observed calls (18,158, or 2.76%), we set the consensus genotype to missing.

We calculated concordance across all common (MAF ≥ 5%) and low frequency (MAF < 5%) markers, where MAFs were assessed over the entire MVP sample. We then calculated concordance between the consensus sequence and each positive control. Concordance was defined as the number of matching called genotypes over the total number of called genotypes. Uncalled markers in either the positive control or the consensus sequence were not included in either the numerator or the denominator of the concordance calculation. We then plotted the concordance distribution for each batch’s positive controls across time.

### Comparing MVP allele frequencies to those from gnomAD and UK Biobank

Genome Aggregation Database (gnomAD) version 2.1 data were downloaded from https://gnomad.broadinstitute.org/downloads. Markers in both gnomAD and MVP were matched on chromosome, start position, end position, reference allele, and alternative allele. For any mismatch, we checked strands and indel notations. Reference and alternative alleles were corrected and frequencies recomputed when strands were flipped. Indels had their genomic coordinates and alleles recoded and harmonized.

UK Biobank summary data were downloaded from https://gbe.stanford.edu. Markers shared between the UK Biobank and MVP were matched using SNP rsIDs. Since information on marker chromosome, genomic positions, reference allele, and alternate allele were not provided in the summary statistics, we were unable to check for swapped alleles. However, we expect variant annotation in MVP and the UK Biobank to be well harmonized as both were genotyped on Axiom^®^ arrays and following the same standard Axiom^®^ marker QC workflow.

For this analysis, European Americans (EA) were defined as samples with greater than 0.9 GBR proportion based on ADMIXTURE results (described below), resulting in a sample size of 311,365. We used PLINK to compute allele frequencies by genetic ancestry subgroup via “--freq” using default filters and quality control parameters.

### Genetic relatedness

We performed additional preprocessing of the MVP dataset before performing the genetic relatedness analysis. We applied standard PLINK 1.9 filters for genotype missingness (>5% removed), MAF (<1% removed), and sample missingness (>5% removed)^11^. We then conducted pairwise relatedness inference using KING 2.1 to identify related pairs^10^. KING explicitly accounts for population structure and is therefore an appropriate algorithm for our sample, which contains diverse genetic ancestry. However, KING is also known to overestimate relatedness in the presence of recent admixture. Therefore, we selected SNPs with low load in PCs 1-3 for a second round of KING as in the UK Biobank^12^.

The first round of KING was run with the command “--related --degree 3” to identify all potential pair of individuals with closer than 3rd degree relatedness. From this result, we excluded all individuals with more than 200 3rd degree relatives and also families with more than 100 members as suspected sample processing artifacts such as low-level sample contamination. Then, a set of unrelated individuals was defined using the largest_independent_vertext_sets() function in the Python version of the igraph tool. Principal component analysis (PCA) was then conducted with the unrelated samples. Only SNPs with greater than 0.01 MAF and less than 0.015 missingness were considered for this PCA. 23 regions of high LD defined in the UK Biobank^18^ were also excluded, and then SNPs were pruned using an R^2^ threshold of 0.1, window of 1000 markers, and step size of 80. In the end, 90,288 SNPs were selected for PCA, which was conducted using PLINK v2.00a2LM with the command “--pca var-wts approx” to obtain variant weights and fast PCA approximation. Low weight SNPs in PC1, PC2, and PC3 were selected by adjusting the absolute weight threshold to keep at least two thirds of the input SNPs which led to 60,118 SNPs being put forward for the next round of KING.

The second round of KING was again conducted with the command “--related --degree 3”. The effect of using SNPs with low weights in PCs 1-3 on the distribution of number of relatives per individual is shown in Figure S10 A-B. We flagged 35 individuals with more than 200 3^rd^ degree relatives (UK Biobank reported 9 individuals with more than 200 3^rd^ degree relatives), as well as all members of two clusters that were tightly interconnected with each other (Supplemental Materials and Methods and Figure S10 C-D, Figure S11).

We defined genetically identical pairs as those having a kinship coefficient of 0.45 or greater (the maximum kinship coefficient output by KING is 0.5). However, given the large number of intentional duplicates samples in our dataset, we only considered genetically identical pairs as monozygotic twin pairs after cross-referencing EHR data as above. Parent-child pairs were defined as those having a kinship coefficient of greater than or equal to 0.19 and less than 0.45 and having less than 0.0025 percent of the genome held with zero alleles identical-by-state (IBS0). Sample pairs with a kinship coefficient greater than or equal to 0.19 and less than 0.45 and IBS0 greater than or equal to 0.0025 were designated full siblings. Any pairs of participants with a kinship coefficient between 0.0884 and 0.19 were inferred to be second-degree or third-degree relatives. To identify potential trios in our sample, we extracted parent-child pairs in which a sample appears twice. We then assessed the kinship coefficient between the other two participants. If the other two participants were not a related pair and consisted of one male and one female, we identified these three samples as a trio.

### Genetic ancestry

For genetic ancestry analysis, we used the same set of markers used for relatedness analysis and applied LD pruning with PLINK (--indep-pairwise 1000 50 0.05), which left us with 50,000 markers.

#### Principal component analysis

For 1000 Genomes Project projection PCA, we merged the MVP dataset with the 1000 Genomes Project Phase 3 reference panel^13^. The 1000 Genomes Project dataset was first filtered to ensure scalable merging with the MVP dataset. Markers with MAF less than 1% and any samples constituting related pairs were removed prior to LD pruning with PLINK using the same parameters as above. We then calculated PCs using the 1000 Genomes Project dataset and projected the MVP samples onto them using EIGENSOFT, version 6.0.1^14^.

We also calculated the PCs on the filtered MVP dataset alone using the FastPCA method from the EIGENSOFT package for within-cohort PCA. For this PCA, we excluded all related individuals, whereas we kept all related individuals in the 1000 Genomes project PCA.

#### ADMIXTURE analysis

In order to quantify ancestry proportions in MVP, we ran the program ADMIXTURE, version 1.3, on the MVP samples in supervised mode using five reference populations from the 1000 Genomes Project dataset as training data^15^. We chose the five reference populations based on their global geographic location to ensure global representativeness. The Yoruba in Ibadan, Nigeria (YRI) samples serve as a proxy for West African ancestry, the Luhya in Webuye, Kenya (LWK) for East African ancestry, the British in England and Scotland (GBR) for European ancestry, the Han Chinese in Beijing, China (CHB) for East Asian ancestry, and the Peruvians from Lima, Peru (PEL) for Native American ancestry (Figure S8C). Participants with more than 80% of their genetic ancestry attributed to one reference population were assigned to that reference. Remaining participants who had greater than 90% of their genetic ancestry derived from two reference populations were assigned to that pair of populations. Any participants not meeting the above two criteria were assigned to a separate subgroup (MVP_OTHER) and were assumed to contain admixture from three or more reference populations.

#### UMAP analysis

We used Uniform Manifold Approximation Projection (UMAP), a dimensionality reduction method that is useful for visualizing both global and local structure in data, to further visualize the genetic ancestry of the MVP cohort. A UMAP embedding was calculated based on the first 10 principal components of unrelated samples using hyperparameters n_neighbors of 15 and min_distance of 0.1, which were suggested by a previous study on UK Biobank data^16^. We then visualized the population structure by projecting subpopulations identified by our ADMIXTURE analysis onto the UMAP embedding.

### GWAS of Height

Height measurements, dates of measurement, dates of birth for each participant were extracted from the VA healthcare system’s EHR. Any height measurement outside the range of 48 to 84 inches was excluded^17^, and inches were converted to meters. Age at measurement was calculated by subtracting the date of birth from the date of height measurement. Individuals younger than 18 or older than 120 years old were excluded. Sex was genetically determined sex by PLINK.

Markers whose genotype missingness was greater than 1%, as well as non-autosomal markers, were removed. Samples whose missingness was over 5% were also excluded. Using the results of the relatedness analysis described below, we also removed all closely related pairs.

After marker and sample filtering, we ran association tests using BOLT-LMM^18^ with sex, age, age-squared and the first 10 PCs as covariates. LD scores were calculated from the 1000 Genomes Project population subsets using ldsc 1.0^19^. Model SNPs were generated using PLINK 2.0 by pruning unrelated samples with an R-squared threshold of 0.2 (--pairwise-indep 1000 50 0.2). Principal components (PCs) were also generated using PLINK 2.0 (--pca approx) on the cohorts that had model SNPs extracted.

We extracted the effect size, direction of effect, and allele for each previously associated marker from the GWAS catalog on March 21, 2019 and then extracted the effects for the markers present in the MVP association analysis. We then scaled the effect values within each study to between 0 and 1 to account for different height units and plotted the previously derived effects against those inferred in MVP.

## Results

### The MVP 1.0 Array

#### Array design and content

The MVP 1.0 array was based on the Applied Biosystems™ Axiom^®^ Biobank Genotyping Array with additional custom content further developed for MVP (Figure 1). The Axiom^®^ Biobank Genotyping Array incorporates multiple content categories that are important for translational medicine research and discovery,^20^ including modules for genome-wide coverage of common European variants, rare coding SNPs and indels, pharmacogenomics markers, expression quantitative trait loci (eQTLs), and loss-of-function markers (further described in Supplemental Materials and Methods). The MVP 1.0-specific modules were mainly SNPs and indels known to be associated with diseases and traits of interest to MVP (especially psychiatric disorders and rheumatoid arthritis), as well as a set of SNPs selected to improve African American imputation performance (Supplemental Materials). In total, 723,305 probesets interrogating 686,682 unique bi-allelic markers (SNPs and indels) based on the GRCh37 genome build were tiled onto the MVP 1.0 array. Among these, 270 are mitochondrial markers, 142 are in the non-pseudoautosomal regions of the Y chromosome, 1,139 are in the pseudoautosomal regions (PAR1 and PAR2) of the X and Y chromosomes, 18,026 are in the non-pseudoautosomal regions of the X chromosome, and the remaining 667,105 markers are autosomal markers (Table S1).

**Figure 1.**
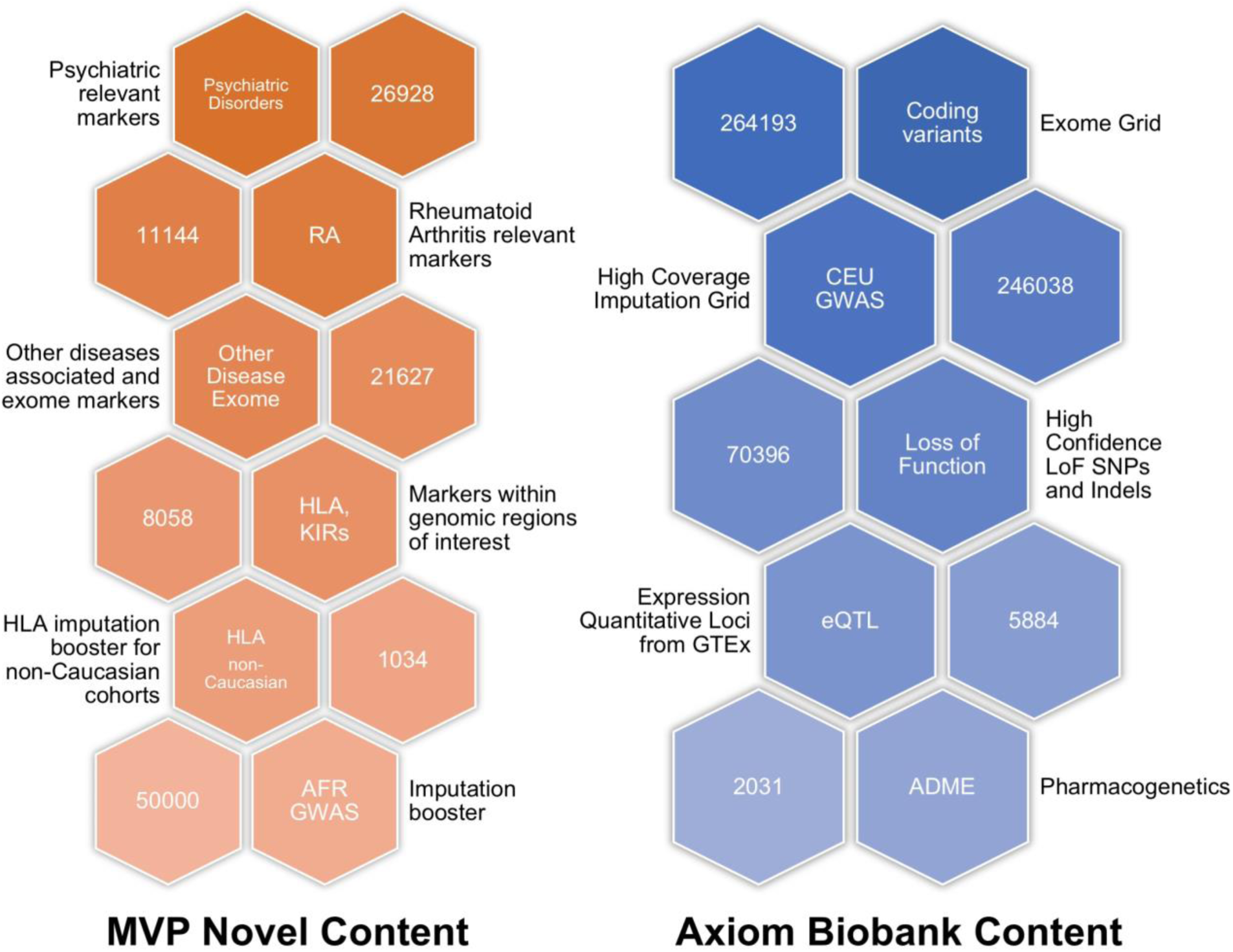
Key MVP 1.0 genotyping array modules. The modules are divided into those shared with the Axiom^®^ Biobank Genotyping Array and those unique to the MVP 1.0 array, along with descriptions and counts of unique markers in each module. Counts represent the number of markers in the module, and markers can be in more than one module.

### MVP 1.0 Genotyping Quality Control and Assessment

#### Assessment of overall genotyping performance

Figure S3 is an overview of the steps taken to ensure high quality genotype data for the MVP cohort. Advanced genotype and sample QC were conducted in addition to the standard Affymetrix good practice guidelines and are described in Materials and Methods and Supplemental Materials and Methods. In addition, we further devised a batch variation correction step to apply to markers that showed significant allele frequency differences between releases (Supplemental Methods and Figure S4, Figure S6A).

We investigated multiple quality control metrics for across and within the two assay vendors. Median Axiom^®^ DQC values for all genotyping batches were greater than 95 for either vendor (Figure S5A). Median QC call rate was also high, exceeding 99% for each genotyping batch (Figure S5 B-C). Overall, sample call rates and other genotype quality control metrics demonstrated high-quality genotype calls for MVP regardless of genotyping vendor (more detail in Supplemental Materials and Methods).

#### Marker and sample QC and selection

The MVP 1.0 array contains a large amount of novel, custom marker content that has not been validated on other arrays. These markers were assayed with more than one probeset, requiring advanced marker QC to determine which probesets for a given marker performed best across all genotyped batches and to remove systematically poor quality probesets. Ultimately, we retained 668,418 markers representing 97.34% of the original markers and included 459,777 samples from a total of 485,856 unique genotyped samples in this data release. As expected, almost 98% of the markers that were previously tested on the Axiom biobank array were associated with a probeset that passed quality control, whereas 77% of the markers in the MVP 1.0 custom modules were associated with a probeset that remained after quality control. Additionally, although sample missingness (the fraction of missing genotype calls per individual; see Supplemental Materials and Methods) was slightly higher for Vendor 1 than for Vendor 2, almost all genotyped samples from both vendors exhibit missingness of less than 5% (Figure S6A).

We also either excluded or quarantined samples that did not meet sample QC criteria. Excluded samples include those expected to be removed by design or for known logistical or data errors. These samples include positive controls, samples with no or multiple unique participant identifiers, and samples in intentional duplicate pairs with the lower call rate. Quarantined samples are those that are temporarily removed from the dataset due to quality concerns. For instance, we investigated 1,149 pairs of samples with high relatedness to discriminate between misidentified intentional duplicates, technical duplicates (controls repeatedly genotyped by vendors), and monozygotic twins. While we confirmed 49 monozygotic twins by cross-referencing with EHR data, the remaining 1,100 unintentional duplicate pairs could not be verified through independent means and were quarantined from data release as potentially mislabeled and will be re-genotyped. We also cross-checked genetically determined sample sex with EHR-reported gender information. Among the 485,856 unique genotyped samples, 2,000 (0.41%) did not have any reported gender information from either the EHR or self-report, and 2,073 (0.43%) of the remaining samples had a genetic sex that was opposite of the reported gender. We quarantined these samples for further analysis and potential re-genotyping (Table S2). The total number of samples that were excluded or quarantined from the current release of MVP genotype data and the reasons for doing so are summarized in Table 1. All quarantined samples removed from the current data release will undergo further quality control validation, be sent back to the vendors for re-genotyping, or will be otherwise verified before being included in subsequent data releases.

**Table 1.** Quarantine and exclusion criteria for MVP samples, and sample count per category.

| Category | Number of samples | Percentage of samples |
| --- | --- | --- |
| **Starting MVP sample set for analysis** | 514,383 |  |
| Intentional duplicate samples | 25,291 |  |
| Uniquely genotyped individuals | 485,856 | 100.00% |
| Samples with call rates below 98.5% | 15,436 | 3.18% |
| Positive control samples | 3,236 | 0.66% |
| Samples with sex misclassification | 1,450 | 0.29% |
| Samples on plates containing 4 or more sex misclassifications | 2,619 | 0.53% |
| Unintentional duplicate samples | 1,149 | 0.23% |
| Samples on plates containing an intentional duplicate with high discordance | 9,975 | 2.05% |
| Samples with high heterozygosity | 248 | 0.05% |
| Samples with no or multiple unique participant identifiers | 71 | 0.01% |
| Intentional duplicate samples with high discordance | 413 | 0.08% |
| Samples with 7 or more “relatives” | 466 | 0.09% |
| **Samples excluded from the dataset** | **28,527** | **5.87%** |
| **Samples quarantined from the dataset** | **31,836** | **6.55%** |
| **Final sample set in current data Release** | **459,777** |  |

Percentages are calculated from the total number of genotyped samples, including positive controls and duplicate samples (514,383). Categories are not mutually exclusive (i.e., a sample can be removed due to more than one category and is counted in each applicable category in the table).

#### Marker missingness and discordance by MAF

We assessed marker missingness in correlation with MAF. Overall, the MAF distribution of MVP 1.0 is highly skewed toward rare variants with 42.89% of markers below 1% MAF and 33.89% below 0.1% (Figure 2A). This result is by design, as the content of the MVP array focuses on markers associated with potential disease phenotypes. We find that MAF is correlated with marker missingness, as shown in Figure 2C and Figure S6B, with lower frequency variants missing in a larger fraction of samples. Despite this trend, missingness among low frequency markers is still relatively low. For example, 87.29% of rare markers (MAF < 0.1%) are missing in less than 5% of genotype calls.

**Figure 2.**
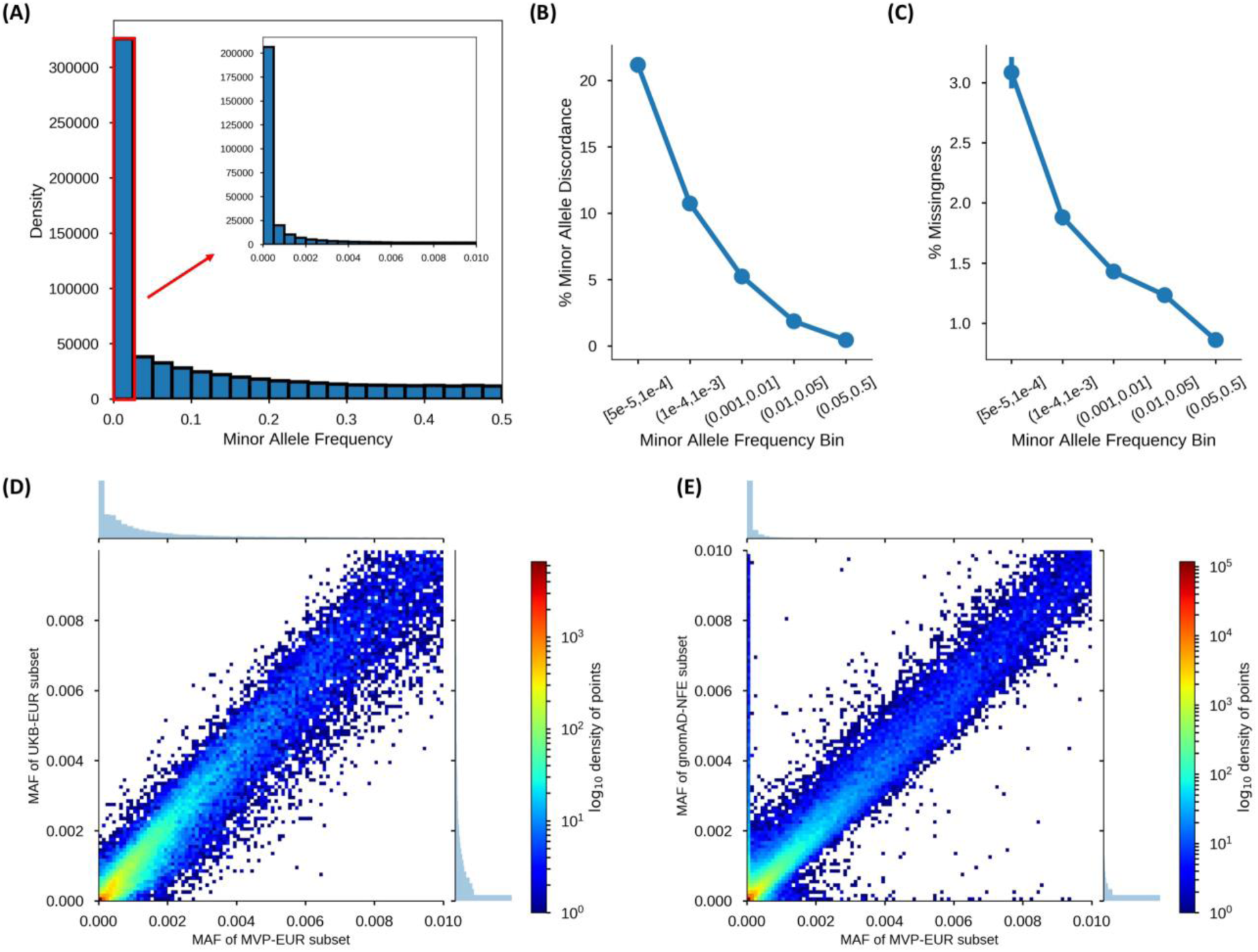
Quality control assessments on the MVP dataset after performing the Advanced Marker Quality Control procedures. (A) MAF distribution after sample QC filtering. The inset diagram shows the distribution for markers with a MAF below 1%. (B) Cumulative fraction of markers for intentional duplicate discordance rates per MAP, separated by MAF bin. (C) Proportion of markers with fraction of missing calls, separated into MAF bins as represented by grayscale color, after sample QC filtering. (D) Comparison of MAFs between the EA subset of MVP (MVP-EUR) and the UK Biobank European subset (UKB-EUR). (D) Comparison of MAFs between MVP-EUR and the non-Finnish European subset of gnomAD (gnomAD-NFE).

Additionally, we examined marker genotype discordance rates across intentional duplicate sample pairs with respect to MAF. Discordance is calculated per minor allele pair (MAP) for each marker, and markers are binned by MAF. We find a correlation between MAF and discordance rate, with lower frequency variants having a higher rate of minor allele discordance (Figure 2B and Figure S6C).

#### Duplicate and positive control samples for continuous quality assessment

Importantly, because we employed two separate vendors for genotyping, we intentionally included 25,291 duplicate samples that were blinded to the vendors for independent assessment of genotype quality. This amounts to a target of 5% of all genotyped samples and is an effort to accurately assess genotyping quality on a continuous basis. Sample concordance among intentional duplicates or positive controls was very high with a median concordance rate greater than 99.8% across all comparisons (Figure S7A).

Assessing concordance in positive control samples also provides valuable information about the consistency and reproducibility of the MVP 1.0 array’s genotypes over time. Along with the MVP samples, 2,064 positive control samples were genotyped on the MVP 1.0 array. As discussed in the Materials and Methods section, we constructed a consensus genotype sequence across 657,459 markers using this panel of positive controls. For markers in the consensus sequence, 543,691 (82.70%) were homozygous, 95,079 (14.46%) were heterozygous, and 18,689 (2.84%) were uncalled. Concordance for each of the 2,064 positive controls samples is defined as the number of markers that agree with the consensus sequence divided by the number of called markers in the consensus sequence. Overall positive control concordance is shown in Figure S7A, and the distributions by batch of concordance values across all positive controls are shown in Figure S7 B-D. The median concordance rate between each positive control sample and the consensus sequence was 99.93% for all markers, 99.89% for common (MAF ≥5%) markers, and 100.00% for low frequency (MAF < 5%) markers. The minimum observed concordance rate between a positive control and the consensus occurs when analyzing common markers, but this concordance rate is still high at 99.05%.

#### Concordance with HapMap samples

To further test concordance and genotyping quality, we genotyped 96 HapMap samples (from Coriell cell lines) on the MVP 1.0 array. 210,630 markers are present in both the MVP 1.0 array and HapMap release 27, and among these markers, 205,647 (97.20%) are classified as recommended (see Supplemental Materials and Methods, Standard marker quality control). When these 205,647 markers were analyzed over the 96 HapMap samples, and when HapMap and Axiom^®^ uncalled genotypes were removed from the numerator and denominator, the sample concordance across all population groups is 99.70% (Table 2). Axiom^®^ sample call rate for recommended markers is 99.85%.

**Table 2.** Concordance rates across 96 HapMap samples genotyped on the MVP 1.0 array.

| Population | Number of samples | Metrics over recommended<sup>a</sup> markers |  | Metrics over all markers |  |
| --- | --- | --- | --- | --- | --- |
|  |  | Average sample concordance (%) | Average sample call rate (%) | Average sample concordance (%) | Average sample call rate (%) |
| ALL | 96 | 99.70 | 99.85 | 99.35 | 99.49 |
| CEU | 28 | 99.70 | 99.85 | 99.34 | 99.47 |
| CHB | 20 | 99.70 | 99.86 | 99.37 | 99.51 |
| JPT | 20 | 99.68 | 99.84 | 99.35 | 99.51 |
| YRI | 28 | 99.71 | 99.86 | 99.34 | 99.49 |

<sup>a</sup> Recommended markers are those that were classified into one of the recommended SNP classes following execution of the Axiom® Best Practices Genotyping workflow for the 96 co-clustered samples.

#### Assessing rare allele genotyping quality

Given the importance of rare markers in clinically-related studies, we evaluated the analytical validity of MVP 1.0 rare markers by observing the concordance of MAFs for rare markers with overlap between MVP 1.0 and either the gnomAD or the UK Biobank (Figure 2 D-E). These databases are large enough for detection of very low MAFs, and agreement of MVP 1.0 marker MAFs with MAFs from these databases provides evidence for the accuracy of MVP 1.0 calls. MAFs were considered to agree when the lower bound of the regression slope’s 95% confidence interval was ≥ 0.9. This value leaves some margin of error for expected differences between the databases in population structure (non-Finnish Europeans vs. European Americans [EA]), technology (genotype arrays vs. exome sequencing), technical processes (batch, user, etc.), and sample size. We used the MVP EA subgroup to benchmark performance because it has a larger sample size which provides better confidence in assessing frequency of rare markers, and has large complementary subgroups in gnomAD and the UK Biobank. We classified markers into three subgroups by MAF: rare variants (< 1%), low frequency variants (1-5%), and common variants (>5%). The EA subgroup yielded 321,290 (48.1%) rare markers, 46,626 (6.97%) low frequency markers, and 300,375 (44.9%) common markers.

From the gnomAD database, we compared the allele frequencies derived from the non-Finnish European subgroup (N = 55,860) of the exome call set. This subgroup provided the largest cohort that was comparable in population structure. In total, a majority of MVP rare variants were found in gnomAD (69%, or 221,374 of 321,290 markers), and we found MAF agreement between MVP and gnomAD with a slope of 0.9290 (95% CI: 0.9002, 0.9578).

From the UK Biobank, we compared allele frequencies derived from the self-reported white British ancestry group (N > 330k). We found MAF agreement as supported by the strong coefficient of determination (R^2^) of 0.9864 and slope of 0.9536 (95%CI: 0.9841, 0.9887) between 46,872 overlapping markers.

While comparison against both sources met the ≥ 0.9 agreement threshold, we observed a small set of about 6000 extremely discrepant markers (defined as having MAF > 0.001 in one database but MAF < 0.001 in the other) between MVP and gnomAD. About 53% of these markers were also present in the UK Biobank. For these discrepant markers, MAFs in the UK Biobank were much closer to MVP MAFs than those in gnomAD, and only one quarter of the overlapping UK Biobank markers retained the “extremely discrepant” label. This is expected and consistent with previous observations that MAFs of MVP and the UK Biobank are in close agreement. The extremely discrepant markers between MVP and gnomAD may be attributed to the gnomAD-exome database having a smaller sample size than the UK Biobank. The lowest MAF limit for MVP’s EA subgroup is 1.6×10^−6^ (1 of 622,730 total alleles), 8.9×10^−6^ (1 of 111,720) for gnomAD’s non-Finnish subgroup, and 1.4×10^−6^ (1 of 674,398) for UK Biobank. At very low frequencies, the absolute difference between rare variants, but not necessarily the relative difference, will be small. A given marker with a MAF of 0.001 in MVP and 0.01 in gnomAD will have an absolute difference of 0.009, but a relative difference of 10-fold. This is a common situation in our pairwise marker comparisons since overlapping marker MAFs are heavily clustered near zero (Figure 2 D-E). This could also explain the relatively higher variance observed in the lower extremes when comparing MVP against gnomAD versus against the UK Biobank. Overall, our results nonetheless show that our rare variant calls are highly consistent and within a reasonable range of agreement with overlapping markers in gnomAD and the UK Biobank. However, it is important to note that precision of very rare variants assayed using SNP chips have been reported to show variable quality^21^. Thus, visual inspection of calls underlying initial association results are always required.

### Population analysis of MVP samples and a test GWAS on height

#### The MVP Cohort

In addition to quality assessment of MVP 1.0 genotyping results, we also performed exploratory analysis of the current population represented in the MVP samples. Based on data from the VistA EHR, the genotyped participants in the MVP cohort have a median age of 65 years at time of enrollment, and 8.33% are female. Although the percentage of female participants is low, reflecting the demographics of the Veteran population, this percentage corresponds to 46,924 female participants in the current release.

Considering the samples that have already been genotyped, the MVP cohort is relatively more diverse than other large biobanks on which data is available. For example, more than 94% of UK Biobank participants self-report as British, Irish, or “any other white background”^4,12^, and the Kaiser RPGEH biobank has 81% of samples reporting as “white, non-Hispanic”. The MVP cohort on the other hand, has 70.9% of participants self-reporting as “white” and “non-Hispanic or Latino” and agrees with United States 2010 census information indicating 63.7% of respondents self-reporting as “White alone” and “Not Hispanic or Latino”^22^.

#### Analysis of relatedness

We examined the degree to which samples in the MVP population are related. Of the approximately 105.70 billion possible MVP sample pairings, 15,384 pairs appeared to be third degree relatives or closer. The number of pairs for each type of relative pair, including trios, is shown in Table S8. Compared with the UK Biobank, this installment of MVP samples has a reduced fraction of related pairs.

#### Analysis of genetic ancestry

Assessing genetic ancestry for genotyped samples is an important tool for many applications, such as correcting for biases caused by population structure, constructing tests for natural selection, and determining disease risk by genetic ancestry, among other tasks^23^. To assess genetic ancestry in our sample, we visualized and then quantitatively assessed genetic ancestry of MVP samples relative to external reference populations.

Runs of homozygosity (ROH) were measured using PLINK with a minimum ROH length of 1,000 Kb. The median total length of ROH is approximately 15.65 Mb, and the median number of blocks per sample is 10. In Figure 3A, we plotted the total length of ROH per individual by genetic ancestry subgroup for the five most common subgroups as defined in the Materials and Methods. MVP_GBR_PEL samples have a wide distribution of total ROH length but also some of the longest total lengths of all samples. Samples with African ancestry or admixed between three or more reference populations (MVP_OTHER) have the shortest total length of ROH per sample. Samples of mainly European ancestry have intermediate total ROH length. The total length of ROH per sample varies depending on the genetic ancestry subgroup.

**Figure 3.**
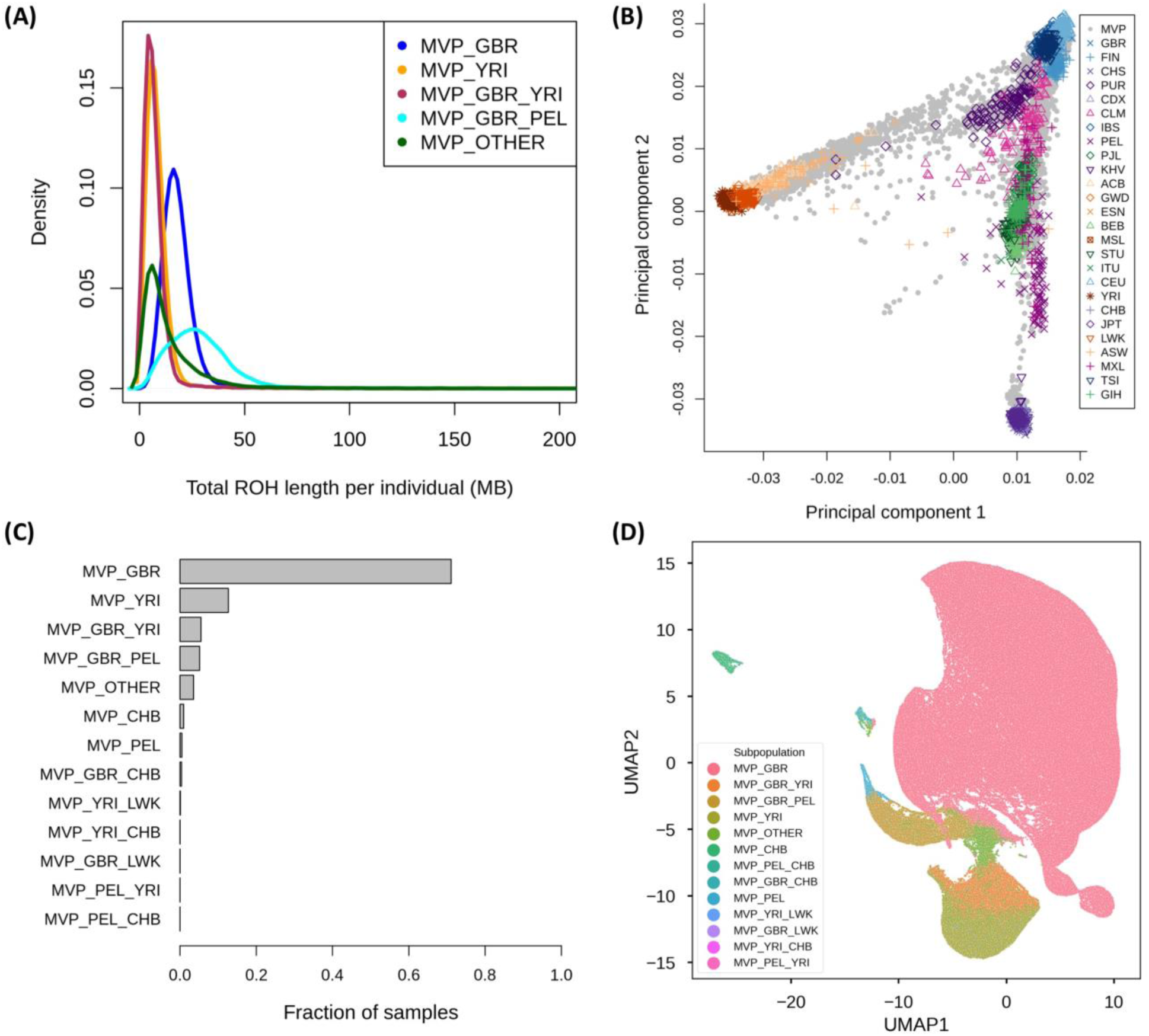
Analysis of genetic ancestry in the MVP dataset. (A) Density plots of the total length of runs of homozygosity (ROH) per individual in each genetic ancestry subgroup. Only the top five most common subgroups are shown. (B) Principal component analysis of the 1000 Genomes Project Phase 3 dataset with MVP samples projected onto principal components 1 and 2. (C) The number of MVP samples in each genetic ancestry subgroup as inferred by ADMIXTURE percentages and our thresholds. Subgroups with no samples are not shown. (D) Visualization of ancestry subgroups using Uniform Manifold Approximation Projection.

We also compared MVP samples to those in the 1000 Genomes Project. We first ran a PCA on the 1000 Genomes Project phase 3 samples and then projected the MVP samples onto these PCs. We find that most MVP samples lie close to reference populations of European origin. In addition, when we performed PCA on MVP samples alone, we found that genetic ancestry subgroups contain more complex intercontinental population structure, with a sizeable fraction of MVP samples exhibiting admixture with respect to African and Asian references samples (Figure 3B, Figure S9).

To assess ancestry proportion for each sample in MVP, we ran the program ADMIXTURE in supervised mode using five 1000 Genomes Project Phase 3 reference populations: Han Chinese in Beijing, China (CHB); British in England and Scotland (GBR); Luhya in Webuye, Kenya (LWK); Peruvians from Lima, Peru (PEL); and Yoruba in Ibadan, Nigeria (YRI)^15^. Most participants have the largest percentage of their genome aligning with the GBR population (Figure S8C). However, a substantial fraction of samples contains a moderate amount of genetic ancestry similar to the YRI reference population. Examples were also found of participants who have almost 100% of their genetic ancestry aligning to each of the five reference populations except for LWK. Using ADMIXTURE analysis results, we grouped the MVP samples into sixteen subgroups and determined the proportion of MVP samples belonging to each (Figure 3C). For example, 326,777 samples have over 80% of their genome aligning with the GBR reference population (MVP_GBR) whereas 58,267 samples have 80% or more of their genome aligning with YRI (MVP_YRI). Excluding samples with more than 80% of their genome aligning to one reference population, 25,295 of the samples have 90% or more of their genome aligning with a combination of GBR and YRI reference populations (MVP_GBR_YRI). Approximately 16,351 samples (MVP_OTHER) have neither 80% of their genome aligning with one reference population nor 90% aligning with a combined pair, indicating substantial admixture between three or more reference populations.

Finally, we visualized the diverse ancestry composition of MVP using a non-parametric dimensionality reduction method called UMAP (Figure 3D). As shown through PCA and ADMIXTURE, the largest cluster corresponds to samples with largely European ancestry. In this visualization, the distance between samples and clusters is not to be directly interpreted as genetic distance. Although there are distinct clusters (such as individuals with Asian ancestry forming a tight cluster within themselves on the top left corner, and another small cluster of likely Polynesians in the middle of the plot), most MVP samples of different ancestries form a large single cluster rather than clusters with distinct breaks. This large cluster shows a continuum of ancestry proportion that transitions from GBR on the top right to different levels of admixture with YRI and PEL proportions. This is in line with a previous report based on 32,000 US individuals in the National Geographic Genographic Project cohort^24^.

#### GWAS of height

To further validate the quality of our genotype data and the utility of MVP 1.0 array, we conducted a GWAS of height in both the EA and African American (AA) MVP subpopulations. EAs were defined as individuals with greater than 90% GBR proportion, and AA were defined as individuals with greater than 60% YRI and less than 40% GBR based on ADMIXTURE results (Figure S8 A-B). Our GWAS of height within EA and AA cohorts showed moderate inflation of λ_GC_=1.12 and λ_GC_=1.13, with pseudo-heritability of 0.396 and 0.378, respectively^19,25,26^, a level comparable to previous association studies in height without genotype imputation^27^.

Of the 822 reported associations with height listed in the GWAS catalog^28^, 230 were present in the MVP EA GWAS, and 209 were present in the MVP AA GWAS. We assessed whether we could replicate effect sizes and direction of effects for markers present in MVP EA and AA GWAS by plotting these against the GWAS catalog effect sizes and direction of effects (Figure 4). For the two subpopulations, the MVP associations perfectly replicated the directions of effect in most markers (two SNPs had near 0 effect size in EA). However, as most GWAS catalog associations are derived from Europeans, the overall correlation across all markers was lower for the AA cohort (r=0.69) compared to the EA cohort (r=0.85).

**Figure 4.**
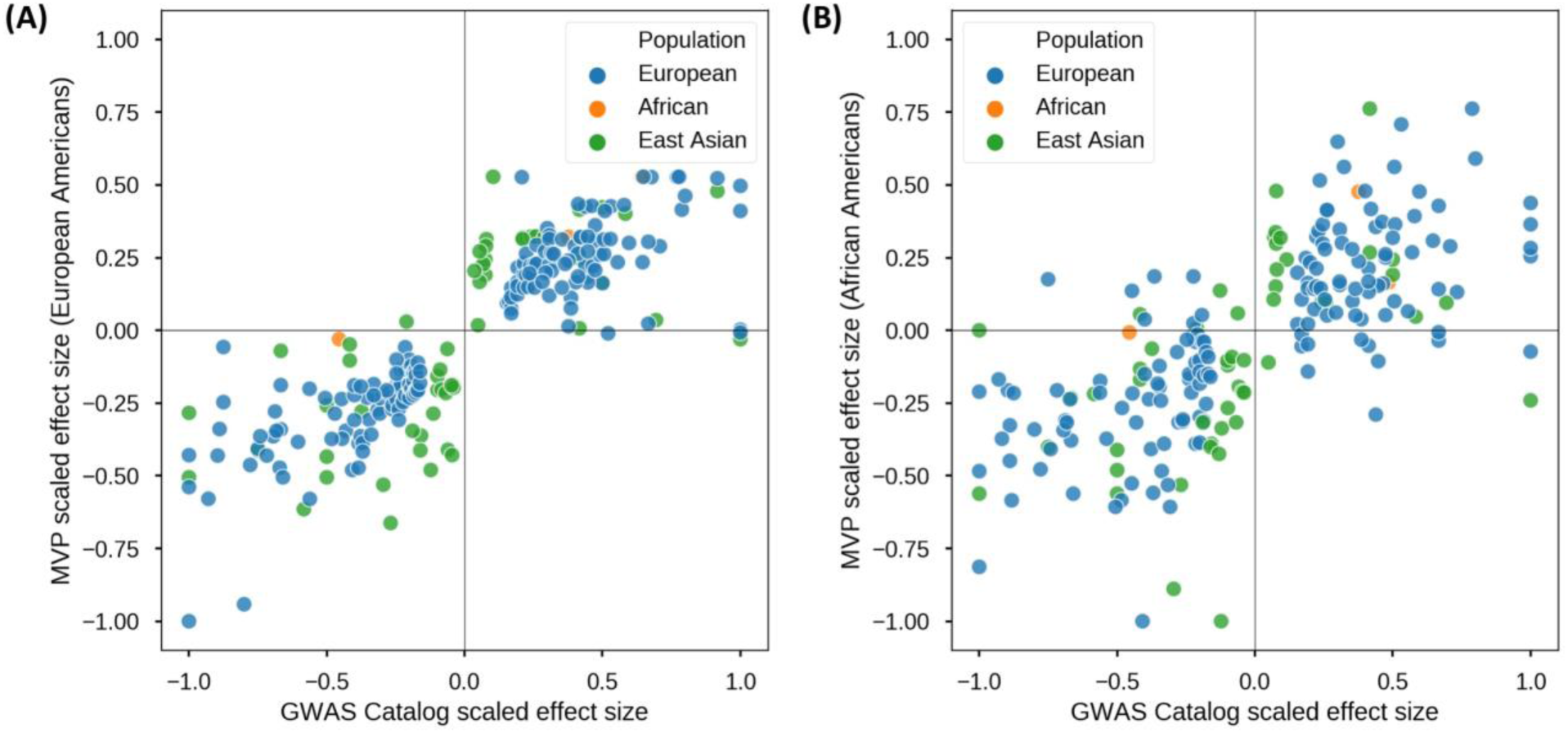
GWAS of height with MVP cohort. (A) Replication of the direction of effect for markers previously associated with height as annotated in the NHGRI-EBI GWAS Catalog in the MVP cohort of non-related European Americans (N=291,609). Color coding denotes the genetic ancestry of the original cohort in which the markers were associated with height. (B) Same as (A) except using the MVP cohort of non-related African Americans (N=73,190).

Overall, we show that the performance of MVP 1.0 and the quality of its genotyping across 459,777 individuals of diverse ethnic background is very consistent and accurate by a variety of metrics.

## Discussion

In this report, we provide an overview of the design of the MVP 1.0 genotyping array, the development of accompanying quality control analyses, and of our initial data exploration of an interim MVP genotyping dataset that consists of nearly 460,000 Veterans. Our results demonstrate that the MVP 1.0 chip and the subsequent QC procedures have addressed notable challenges characteristic of large projects with individuals of diverse genetic background, and that the resulting genotype calls is of high-quality akin to other projects similar in scope. By using a single chip and unified quality control across the diverse cohort, we aimed to minimize batch effects between different ancestries and provide an initial genome-wide scan before whole genome sequenced samples become available.

### Addressing the challenges of MVP

MVP’s large, diverse and still-growing cohort poses numerous challenges for designing genotyping procedures and their subsequent quality assessment/quality control protocols. Genotyping large and ethnically diverse cohorts along with clinically relevant markers is even more challenging due to the finite number of probesets that can fit on a single array. However, using different arrays for different ethnic groups can also exacerbate the differences between these groups and lead to batch effects.

To address the limitations of array-based genotyping in diverse cohorts, we carefully selected array content to maximize clinical utility while at the same time ensuring both broad coverage of variants and robust imputation capabilities across different ethnic groups. We also developed comprehensive quality controls for markers and samples both before and after genotyping, including: intentional duplication of ∼5% randomly selected samples over time, blinded to assay technicians, to detect and mitigate batch variation; assessment of genotyping concordance using positive control samples and HapMap samples (Figure S7A, Table 2); comparing MVP 1.0 MAFs to those in gnomAD and the UK Biobank (Figure 2); and conducting a GWAS of height to replicate previously reported results (Figure 4). Overall, we retained and released 459,777 samples and 668,418 markers after QC for the initial release of data. Although QC metrics vary slightly over time and genotyping vendors, the final genotyped sample set show consistently high call rates (98.5%) and genotype concordance over intentional duplicates (99.8%) both within and between vendors and over time. Furthermore, marker concordance is also high even for rare markers. Additionally, genotype concordance, MAF, and GWAS association results are generally in strong agreement with external or previously reported results. These results indicate that the design of the MVP 1.0 array and the associated quality control and assessment procedures provide a robust, reliable method for both genotyping common, low-frequency, and rare variants in a large, ethnically diverse cohorts.

Challenges remain, however, and the MVP 1.0 array has several limitations. Notably, although concordance rates were high, our results demonstrate that low-frequency and rare variants are still more difficult than common variants to genotype accurately using the MVP 1.0 array. Additionally, while we added markers to MVP 1.0 to increase coverage for African Americans, we lack boosters for other ethnic groups, such as Asian and Native American populations, which currently comprise smaller but growing proportions in the MVP population.

### The MVP dataset is ethnically and genetically diverse

Our exploratory analysis indicates that the MVP dataset and samples offer unique value for disease research. One particularly valuable aspect of the MVP dataset is the ethnic diversity it encompasses. Genetic ancestry analysis suggests that the MVP dataset contains sub-populations with both homogeneous and admixed genetic ancestry from multiple global populations. The largest sub-population corresponds to 71% samples of mostly European descent, with the remaining samples showing substantial African, East Asian, and Native American ancestry.

Since MVP recruits participants from United States Veterans who receive care at VA hospitals, the demographics of the MVP dataset diverge from those of the United States population. Approximately 8.5% of MVP samples are female, which is similar to the fraction of women in the Veteran population^29^. MVP participants are also substantially older than the United States population with a median age of 68 as opposed to 37.9 years^30^. However, the demographics of MVP may change with increasing use of the VA by more recent Veterans who have completed their service. The proportion of female Veterans is projected to continuously grow and nearly double to 16.5% by 2043^29^. Meanwhile, the proportion of Veterans from minority populations is expected to increase by approximately 50% over the same time period^29^. Thus, the VA and MVP is in a unique position for further inclusion of participants from diverse backgrounds.

### The MVP dataset is an invaluable disease research resource

MVP has several unique features that make it an invaluable resource for human disease research. As evidence of the general utility of this resource, initial reports using an earlier tranche of ∼300,000 genotyped participants have reported substantial new findings regarding the genetics of blood lipids, a major cardiovascular risk factor^31^. Not only is MVP ideal for studying the burden of chronic disease, which increases with age, many of the clinical records in its EHR span several decades, allowing for robust longitudinal analysis. This is possible as patients using the VA health services do not lose coverage even after changing employers or residence. Additionally, MVP provides an opportunity to study diseases disproportionately affecting US veterans, such as PTSD^32^, alcohol and substance abuse disorders^33^, as well as other deployment-related conditions and their impact on human health.

In conclusion, the high-quality genotype data generated using the MVP 1.0 array provides a valuable resource for researchers investigating the effect of both rare and common genetic variants within MVP. This quality-controlled genotype data as well as the results from genetic ancestry and relatedness analyses are made available to all approved researchers. The genotype data can be linked to the full EHR of participants, often covering decades of care provided by the VA. MVP is a continuously expanding research cohort made available by participants with diverse backgrounds and altruistic intentions to support research that will benefit their fellow Veterans and others.

## Supplemental Data

Supplemental Data include 11 Figures and 6 Tables.

## Author Contribution

HH-Z, YS, ML, BRG, SJ, NS, TW, AL, PD Performed analysis, TW, LR, JS, TJO, SP Designed array, YS, PH, PD, PK, SP Built Data and software pipelines platforms and optimized them, DH, MB Performed wet lab assays for blood processing and isolating DNA, HH-Z, YS, ML, BRG, SJ, NS, TLA, JH, CP, JM, SM, GDH, RP, JC, JMG, JG, CJO, ERH, HZ, TJO, PST, SP are members of MVP genomic working group, SP conceived and supervised the work.

## Acknowledgments

This work was funded by the VA Office of Research and Development and the VA Special Fellowship in Medical Informatics. We would like to thank all Veteran participants in the MVP for donating their samples, information, and time to this project.

In addition, We acknowledge the Million Veteran Program (MVP) Consortium: MVP Executive Committee, J. Michael Gaziano, MD, MPH (VA Boston Healthcare System, Boston MA), Rachel Ramoni, DMD, ScD (Office of Research and Development, Veterans Affairs Central Office; Washington, DC), Jim Breeling, MD (ex-officio) (Office of Research and Development, Veterans Affairs Central Office; Washington, DC), Kyong-Mi Chang, MD (Philadelphia Veterans Affairs Medical Center; Philadelphia, PA), Grant Huang, PhD (Office of Research and Development, Veterans Affairs Central Office; Washington, DC), Sumitra Muralidhar, PhD (Office of Research and Development, Veterans Affairs Central Office; Washington, DC), Christopher J. O’Donnell, MD, MPH (VA Boston Healthcare System, Boston, MA), Philip S. Tsao, PhD (VA Palo Alto Health Care System; Palo Alto, CA), MVP Program Office, Sumitra Muralidhar, PhD (Office of Research and Development, Veterans Affairs Central Office; Washington, DC), Jennifer Moser, PhD (Office of Research and Development, Veterans Affairs Central Office; Washington, DC), MVP Recruitment/Enrollment, Recruitment/Enrollment Director/Deputy Director, Boston – Stacey B. Whitbourne, PhD; MVP Coordinating Centers, Clinical Epidemiology Research Center (CERC), West Haven – John Concato, MD, MPH (VA Connecticut HealthCare System; West Haven, CT), Cooperative Studies Program Clinical Research Pharmacy Coordinating Center, Albuquerque – Stuart Warren, JD, Pharm D; Dean P. Argyres, MS (Albuquerque VA Medical Center; Albuquerque, NM), MVP Information Center, Canandaigua – Brady Stephens, MS, Core Biorepository, Boston – Mary T. Brophy MD, MPH; Donald E. Humphries, PhD (VA Boston Healthcare System, Boston, MA), Data Operations/Analytics, Boston – Xuan-Mai T. Nguyen, PhD (VA Boston Healthcare System, Boston, MA), MVP Science – Christopher J. O’Donnell, MD, MPH; Genomics - Saiju Pyarajan PhD; Philip S. Tsao, PhD (VA Boston Healthcare System, Boston, MA & VA Palo Alto Health Care System; Palo Alto, CA), Phenomics – Kelly Cho, MPH, PhD (VA Boston Healthcare System, Boston, MA), Data and Computational Sciences – Saiju Pyarajan, PhD (VA Boston Healthcare System, Boston, MA), MVP Local Site Investigators: Atlanta VA Medical Center (Peter Wilson, MD); Bay Pines VA Healthcare System (Rachel McArdle, PhD), Birmingham VA Medical Center (Louis Dellitalia, MD), Cincinnati VA Medical Center (John Harley, MD), Clement J. Zablocki VA Medical Center (Jeffrey Whittle, MD), Durham VA Medical Center (Jean Beckham, PhD), Edith Nourse Rogers Memorial Veterans Hospital (John Wells, PhD), Edward Hines, Jr. VA Medical Center (Salvador Gutierrez, MD), Fayetteville VA Medical Center (Gretchen Gibson, DDS), VA Health Care Upstate New York (Laurence Kaminsky, PhD), New Mexico VA Health Care System (Gerardo Villareal, MD), VA Boston Healthcare System (Scott Kinlay, PhD), VA Western New York Healthcare System (Junzhe Xu, MD), Ralph H. Johnson VA Medical Center (Mark Hamner, MD), Wm. Jennings Bryan Dorn VA Medical Center (Kathlyn Sue Haddock, PhD), VA North Texas Health Care System (Sujata Bhushan, MD), Hampton VA Medical Center (Pran Iruvanti, PhD), Hunter Holmes McGuire VA Medical Center (Michael Godschalk, MD), Iowa City VA Health Care System (Zuhair Ballas, MD), Jack C. Montgomery VA Medical Center (Malcolm Buford, MD), James A. Haley Veterans’ Hospital (Stephen Mastorides, MD), Louisville VA Medical Center (Jon Klein, MD), Manchester VA Medical Center (Nora Ratcliffe, MD), Miami VA Health Care System (Hermes Florez, MD), Michael E. DeBakey VA Medical Center (Alan Swann, MD), Minneapolis VA Health Care System (Maureen Murdoch, MD), N. FL/S. GA Veterans Health System (Peruvemba Sriram, MD), Northport VA Medical Center (Shing Shing Yeh, MD), Overton Brooks VA Medical Center (Ronald Washburn, MD), Philadelphia VA Medical Center (Darshana Jhala, MD), Phoenix VA Health Care System (Samuel Aguayo, MD), Portland VA Medical Center (David Cohen, MD), Providence VA Medical Center (Satish Sharma, MD), Richard Roudebush VA Medical Center (John Callaghan, MD), Salem VA Medical Center (Kris Ann Oursler, MD), San Francisco VA Health Care System (Mary Whooley, MD), South Texas Veterans Health Care System (Sunil Ahuja, MD), Southeast Louisiana Veterans Health Care System (Amparo Gutierrez, MD), Southern Arizona VA Health Care System (Ronald Schifman, MD), Sioux Falls VA Health Care System (Jennifer Greco, MD), St. Louis VA Health Care System (Michael Rauchman, MD), Syracuse VA Medical Center (Richard Servatius, PhD), VA Eastern Kansas Health Care System (Mary Oehlert, PhD), VA Greater Los Angeles Health Care System (Agnes Wallbom, MD), VA Loma Linda Healthcare System (Ronald Fernando, MD), VA Long Beach Healthcare System (Timothy Morgan, MD), VA Maine Healthcare System (Todd Stapley, DO), VA New York Harbor Healthcare System (Scott Sherman, MD), VA Pacific Islands Health Care System (Gwenevere Anderson, RN), VA Palo Alto Health Care System (Philip Tsao, PhD), VA Pittsburgh Health Care System (Elif Sonel, MD), VA Puget Sound Health Care System (Edward Boyko, MD), VA Salt Lake City Health Care System (Laurence Meyer, MD), VA San Diego Healthcare System (Samir Gupta, MD), VA Southern Nevada Healthcare System (Joseph Fayad, MD), VA Tennessee Valley Healthcare System (Adriana Hung, MD), Washington, DC VA Medical Center (Jack Lichy, MD, PhD), W.G. (Bill) Hefner VA Medical Center (Robin Hurley, MD), White River Junction VA Medical Center (Brooks hRobey, MD), William S. Middleton Memorial Veterans Hospital (Robert Striker, MD).

The content of this manuscript does not represent the views of the Department of Veterans Affairs or the United States Government.

## Web Resources

**gnomAD**: https://storage.googleapis.com/gnomad-public/release/2.1/vcf/exomes/gnomad.exomes.r2.1.sites.chr*.vcf.bgz

UK Biobank: https://github.com/rivas-lab/public-resources/blob/master/uk_biobank/variant_filter_table.tsv

