## Supplemental materials for "Measuring genetic variation in the multi-ethnic Million Veteran Program (MVP)"

### 20191221_MVP_GENOTYPING SUPPLEMENTAL FILE

DECEMBER 21, 2019

VETERAN AFFAIRS

A

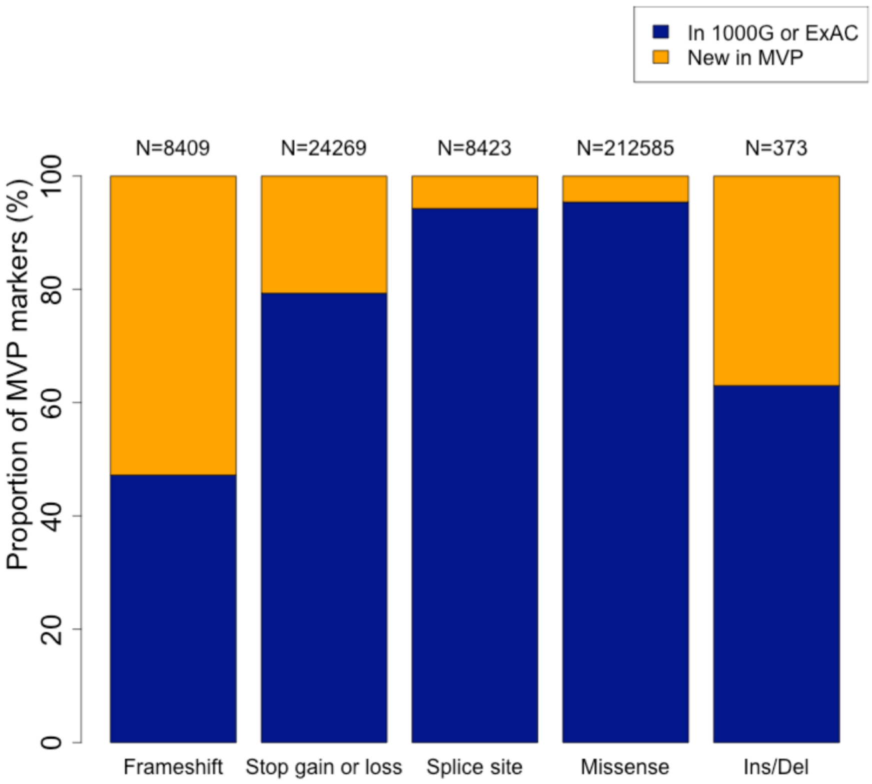


**Figure S1. Comparison of MVP 1.0 markers to those in other databases.** The distribution of MVP 1.0 Array markers by their functional annotation and whether they overlap with existing genetic variation databases, the 1000 Genomes Project and the ExAC database.

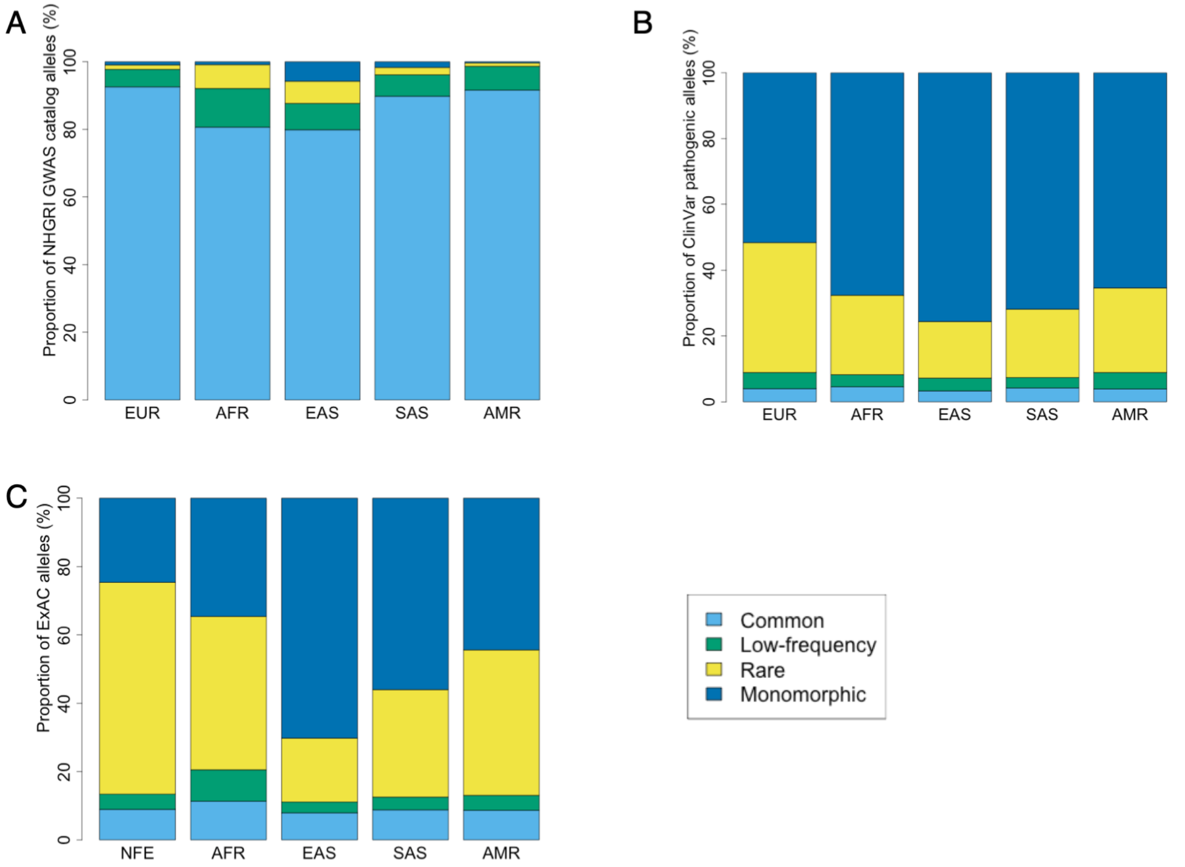


**Figure S2. MVP 1.0 Array marker overlap with databases of functional and disease-related variants.** Distribution of markers overlapping with (A) the NHGRI-EBI GWAS catalog; (B) the ClinVar database; and (C) the ExAC database by MAF bins and ancestral population groups. MAFs and ancestral population definitions were extracted for (A) and (B) from the 1000 Genomes Project and for (C) from the ExAC database. Common variants are markers with MAF ≥ 5%; low-frequency variants are markers with 1% ≤ MAF < 5%; rare variants are markers with MAF < 1%; and monomorphic variants are markers with MAF = 0%. (EUR: European, NFE: Non-Finnish European, AFR: African, EAS: East Asian, SAS: South Asian, AMR: Admixed American)

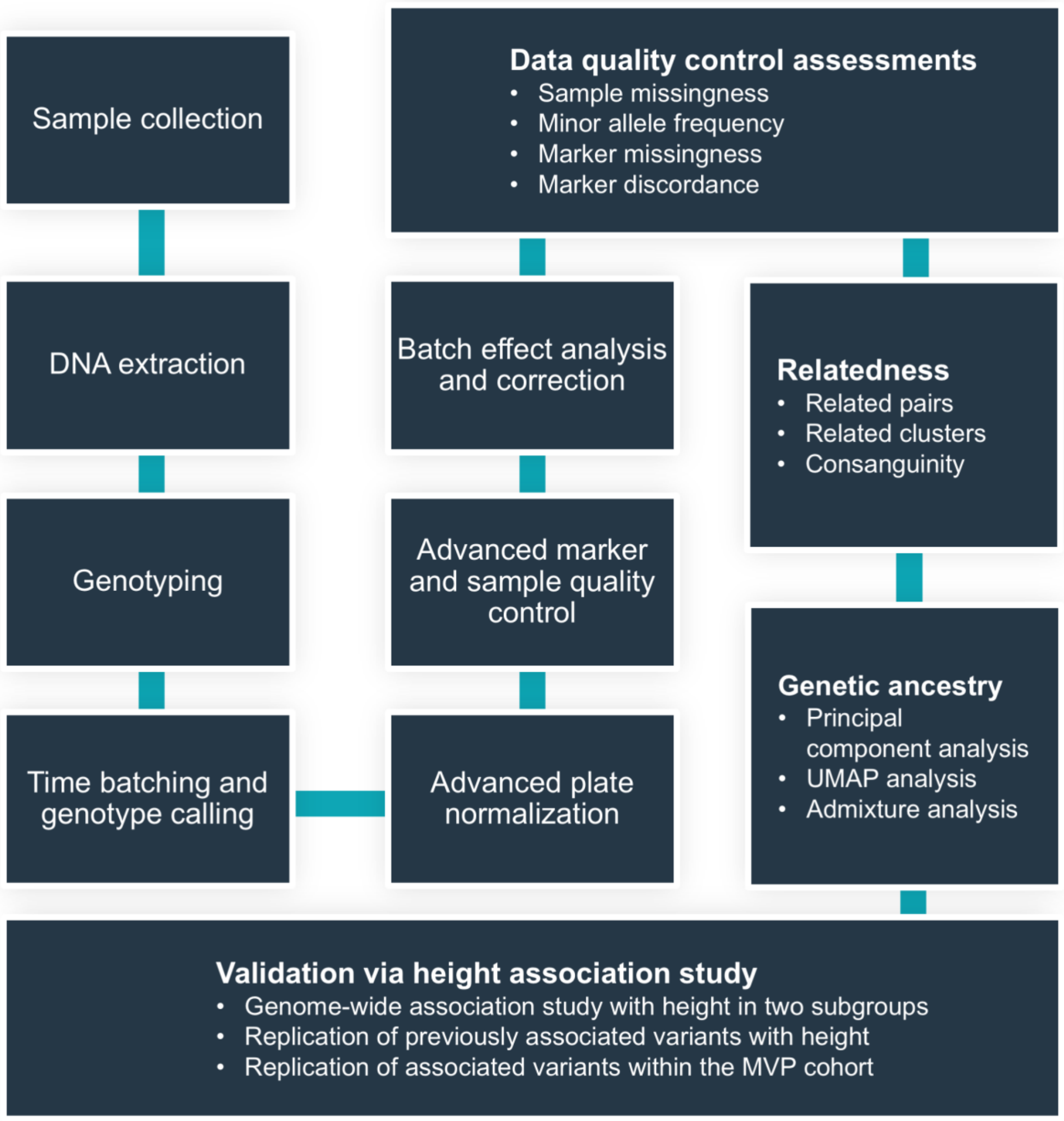


**Figure S3. The overall genotyping quality control and MVP data analysis procedure.** Overview of the steps of the genotype calling, quality control, data exploration, and final validation processes for the MVP dataset.

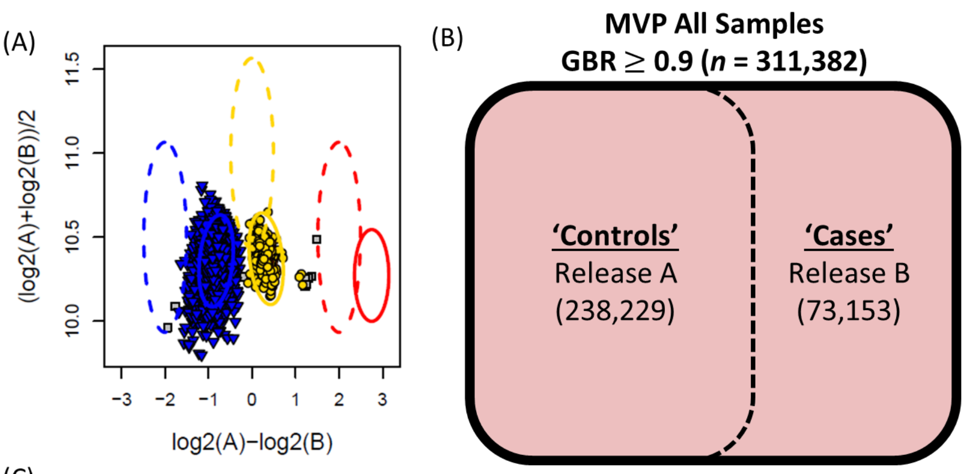


(C)

| Total (N) Batch-SNP Combinations Set to 'No Call' | Filter Added | Significant Associations in GWAS* |
|---|---|---|
| 50,319 | Fisher's Exact Test Only | 73 |
| 144,110 | +No Minor Hom. / HWE | 66 |
| 208,724 | +Low CR/FLD | 48 |
| 231,525 | +Cluster Number | 48 |

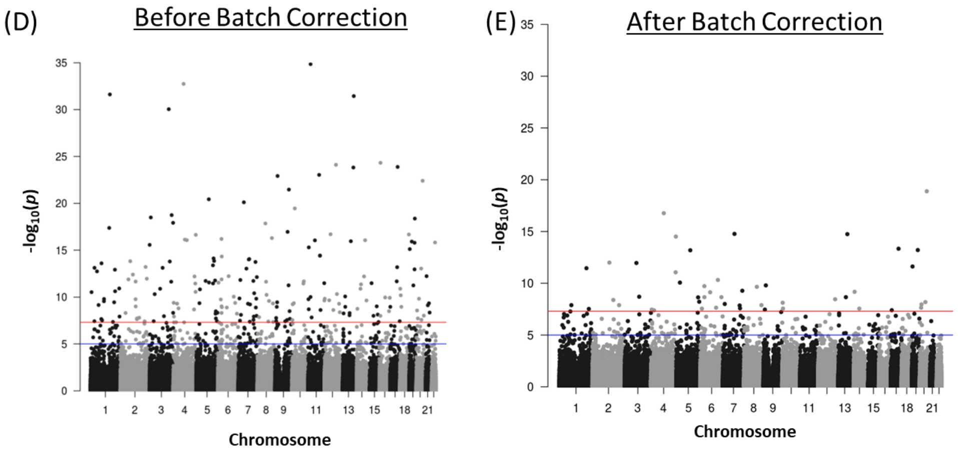


**Figure S4**. **Batch variation correction.** (A) Example of a mis-clustered probeset in a batch detected using Fisher's Exact Test. (B) The GWAS experimental design used to detect batch effects across releases. Release A samples were designated "Controls," and Release B samples were designated "Cases." (C) Filters introduced in addition to Fisher's Exact Test, and the number of batch-SNP combinations removed and significant GWAS associations remaining after each step. (D) Manhattan plot showing 204 significant associations before batch correction. (E) Manhattan plot showing 48 remaining significant associations after application of batch correction procedure.

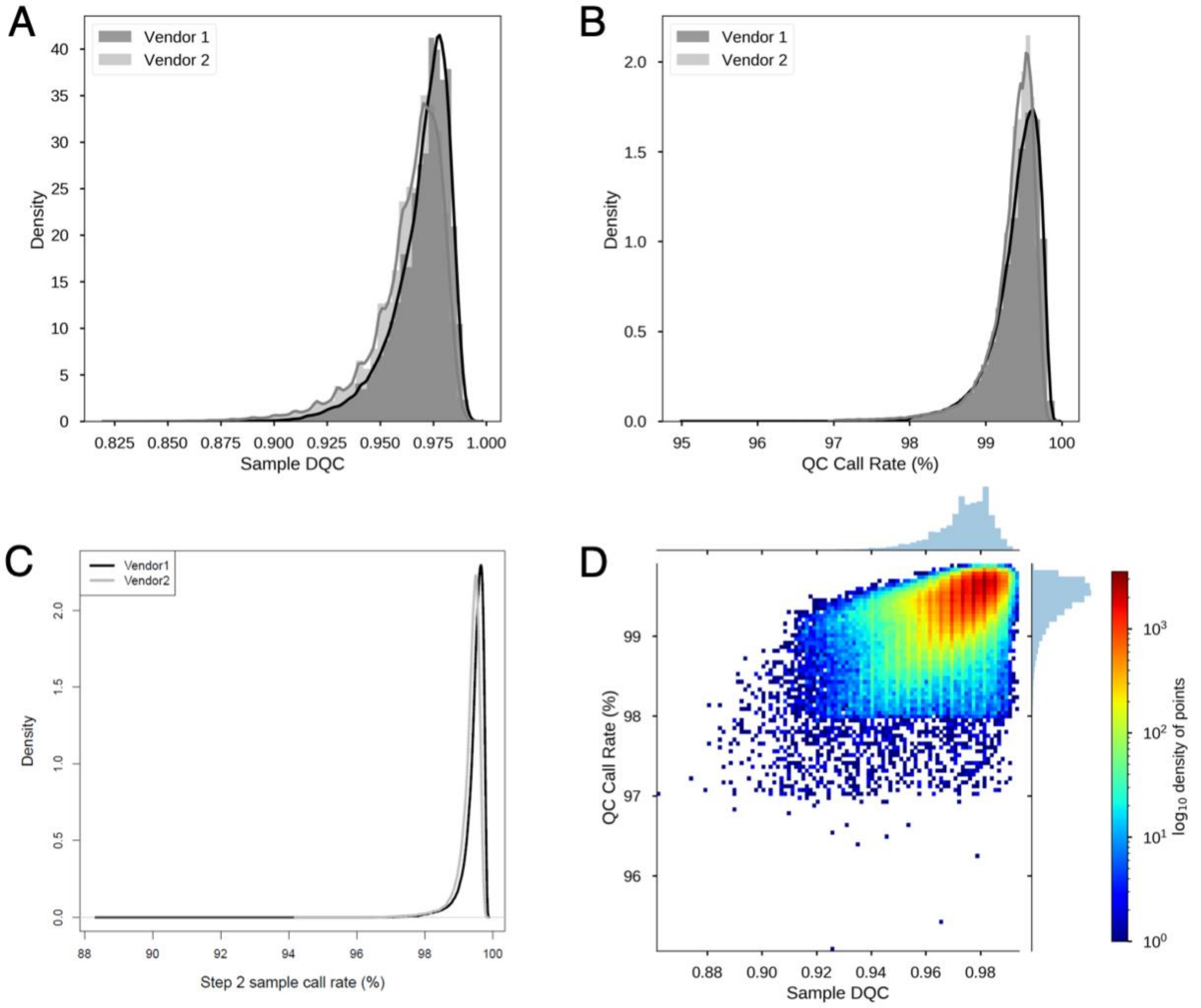


**Figure S5. Axiom® genotyping DQC, sample QC call rate and sample call rate metrics.** Metrics were reported directly from vendors, and distributions for each metric are for all samples genotyped before sample QC filtering. (A) Density curves of the distributions of the dish QC (DQC) metric by genotyping vendor. (B) Density curves of the distribution of sample QC call rate by genotyping vendor. Sample QC call rate is referred to as Step 1 call rate in the Axiom® Genotyping Solution Data Analysis Guide. (C) Density curves of the distribution of the sample call rate over all MVP 1.0 array probesets by genotyping vendor after re-batching and recalling genotypes at the VA. Sample call rate is referred to as Step 2 call rate in the Axiom® Genotyping Solution Data Analysis Guide. (D) Density plot of sample QC call rate against the DQC metric before sample quality control filtering. Color density is scaled by the log of the number of samples, with red indicating higher density and blue indicating lower density.

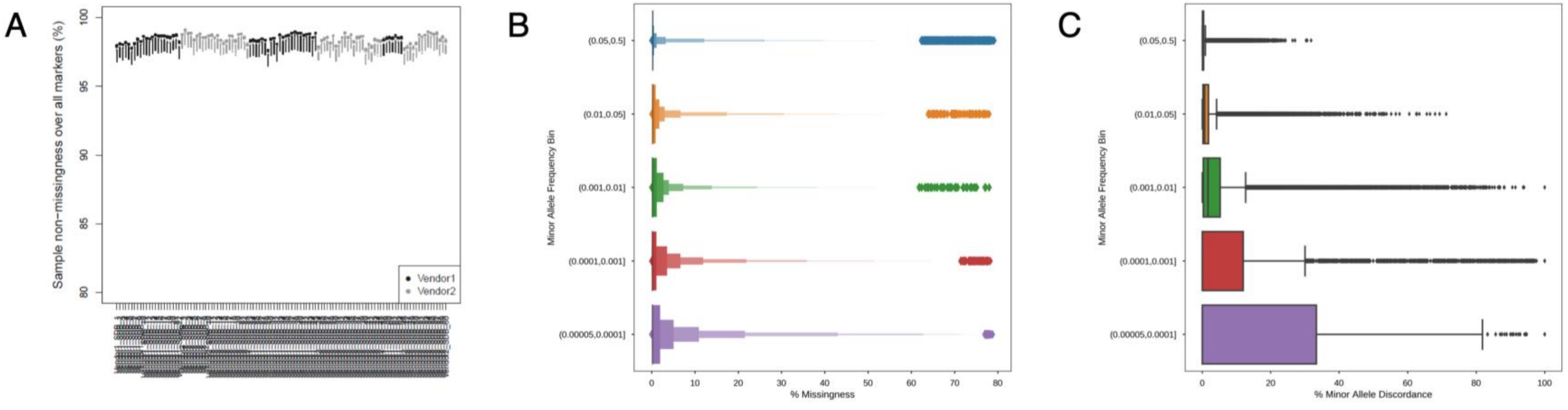


**Figure S6. Assessments of sample and marker missingness.** (A) Sample non-missingness (i.e. 1 – missingness) for each batch. The X-axis conains identifiers for genotyping batches in sequential time order. (B) Percent marker missingness startified by MAF bin. (C) Minor allele discordance by MAF bin.

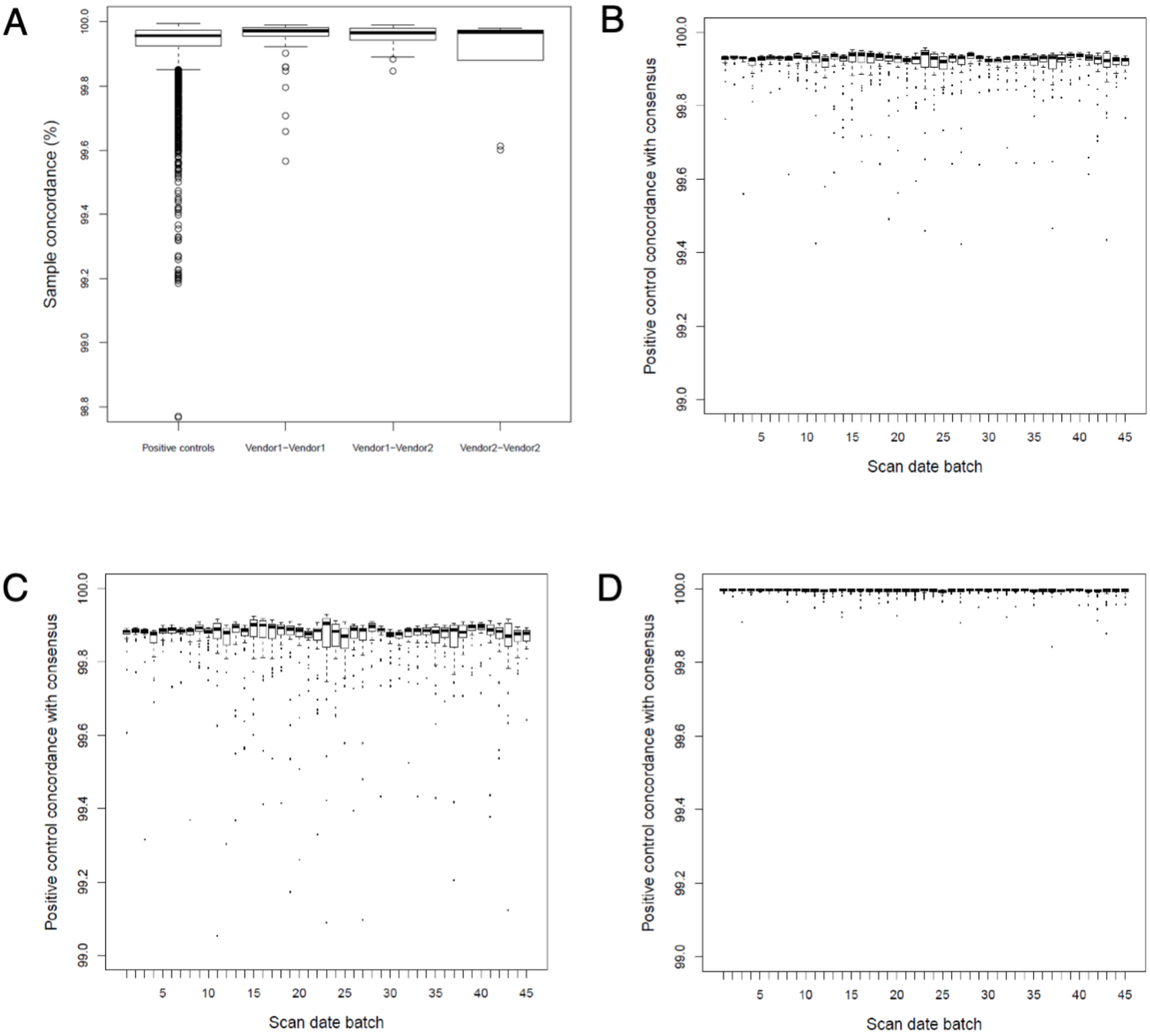


**Figure S7. Sample and marker concordance.** (A) Sample concordance over the marker set after the advanced marker QC procedure for pairs of positive controls and pairs of intentional duplicates genotyped both within and between vendors. Positive controls are only from Vendor 2, since Vendor 1 did not include positive controls on genotyped plates. Distributions are shown for a randomly selected 5,000 pairs for each category. (B-D) Marker concordance for (B) all markers, (C) common (MAF ≥ 5%) markers, and D) low-frequency (MAF < 5%) markers between positive control samples and the consensus positive control genotyping sequence, grouped by batch. Batches are ordered by time.

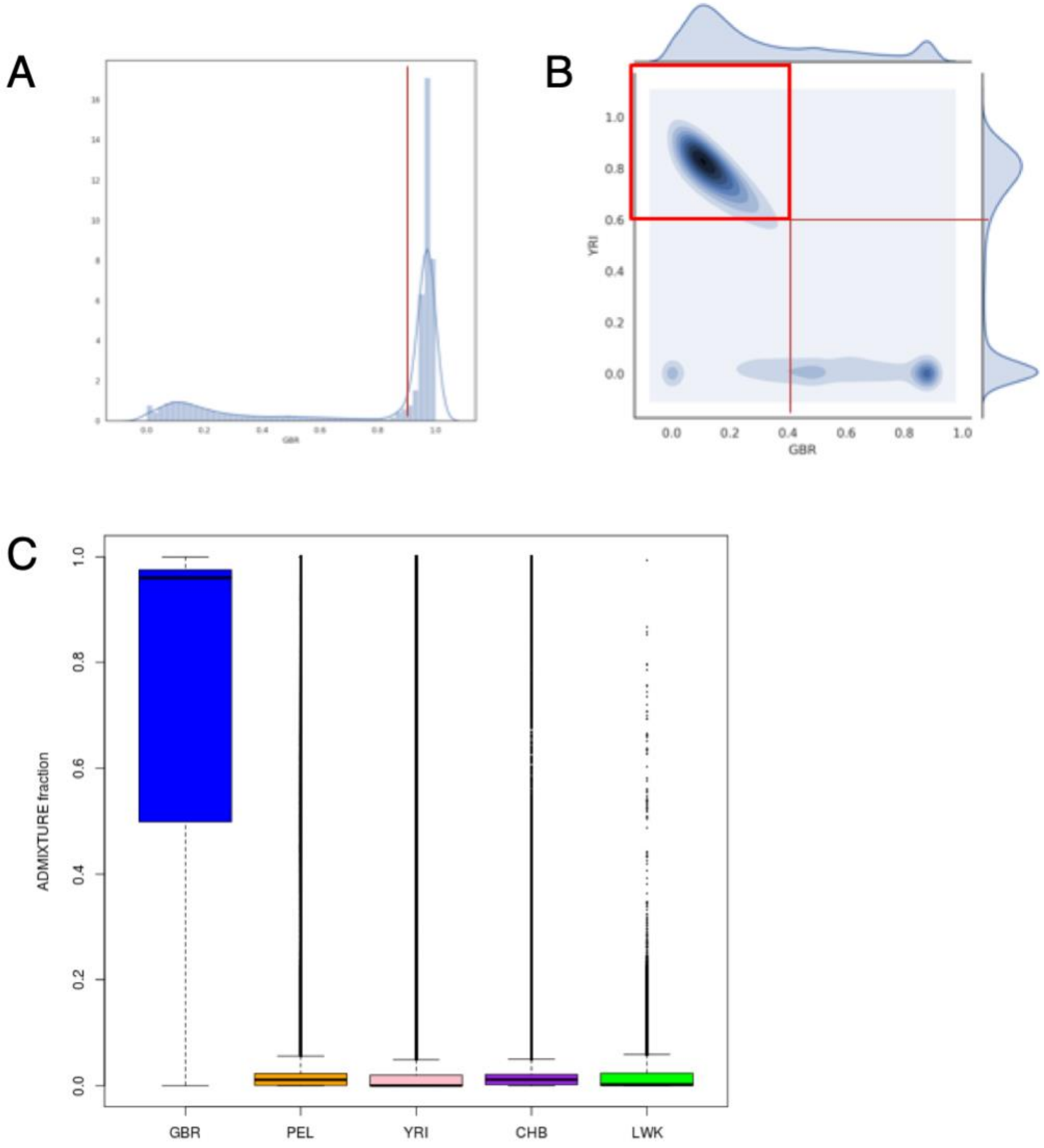


**Figure S8. Genetic ancestry analysis of the MVP cohort using ADMIXTURE.** (A) Distribution of British in England and Scotland (GBR) proportion in all MVP individuals. Individuals with more than 0.9 GBR proportion were defined as European American. (B) Kernel density estimates of Yoruba in Ibadan, Nigeria (YRI) and GBR proportion within individuals with less than 0.9 GBR proportion (i.e. non-European individuals). Individuals with more than 0.6 YRI and less than 0.4 GBR proportion were defined as African American. (C) Distribution of ADMIXTURE proportions for each of the five 1000 Genomes Project reference populations, sorted by average fraction, across all MVP samples. Reference population abbreviations: British in England and Scotland (GBR); Yoruba in Ibadan, Nigeria (YRI); Peruvians from Lima, Peru (PEL); Han Chinese in Beijing, China (CHB); and Luhya in Webuye, Kenya (LWK).

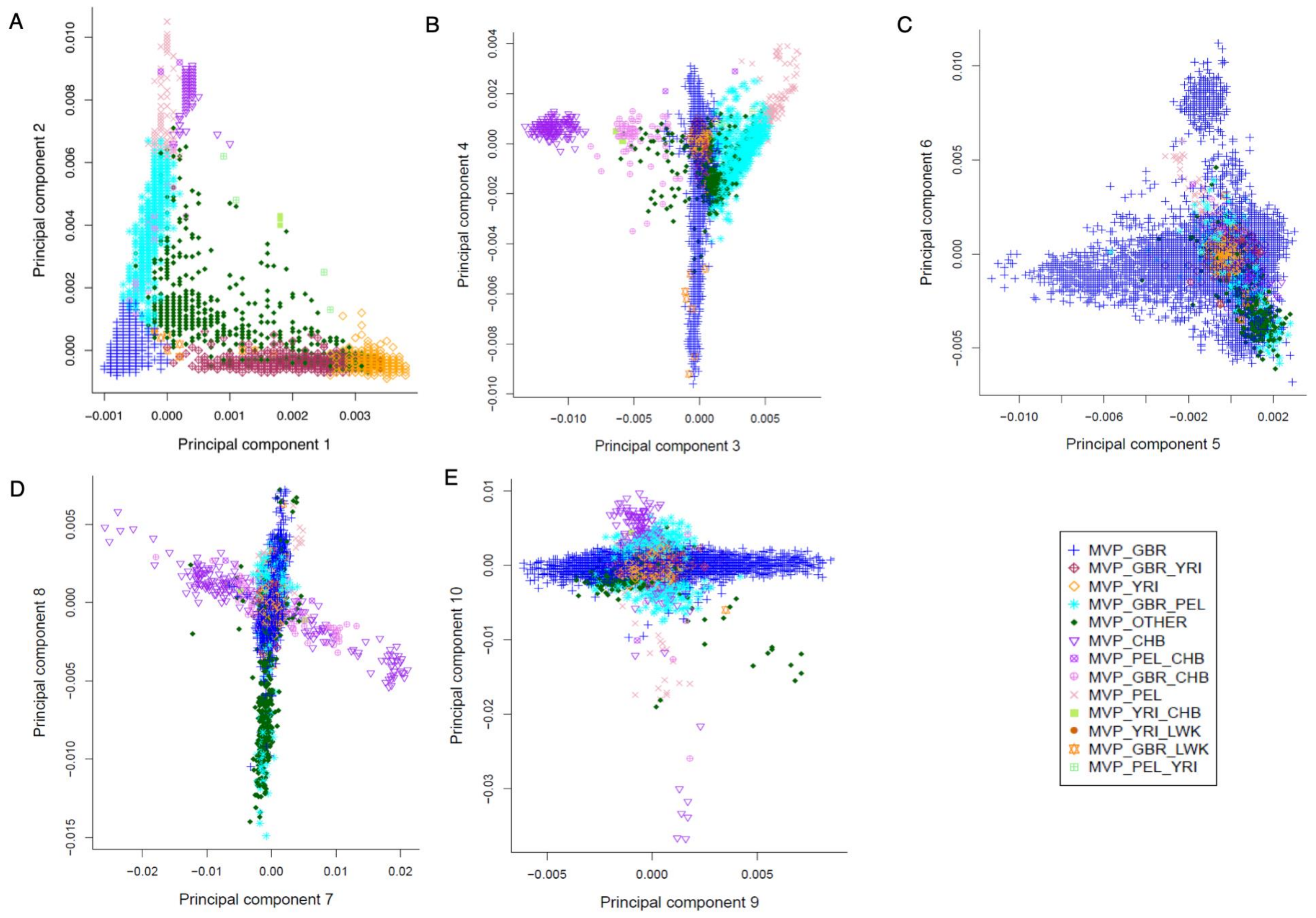


**Figure S9. Ancestry Principal Component Analysis (PCA) of the MVP Cohort.** Colors represent genetic ancestry subgroups as defined by ADMIXTURE analysis. (A) Principal components (PCs) 1 and 2 of the MVP cohort mimic patterns seen in PCA of the 1000 Genomes Project population. (B) PC3 splits samples aligning with the East Asian reference population (MVP_CHB) from the Native American reference population (MVP_PEL). (C) PC 6 reveals a separate cluster of samples with European Ancestry (MVP_GBR) that splits out distinctively from the main cluster of samples in this subgroup. (D) PC 7 reveals substantial diversity in the MVP_CHB subgroup, whereas PC 8 separates MVP_GBR participants from those with admixture from three or more global reference populations (MVP_OTHER). PC 8 also reveals substantial diversity within participants admixed between European and American ancestries (MVP_GBR_PEL). (E) PC 10 reveals a gradient between MVP_GBR and MVP_PEL samples, with MVP_GBR_PEL samples lying between the two clusters.

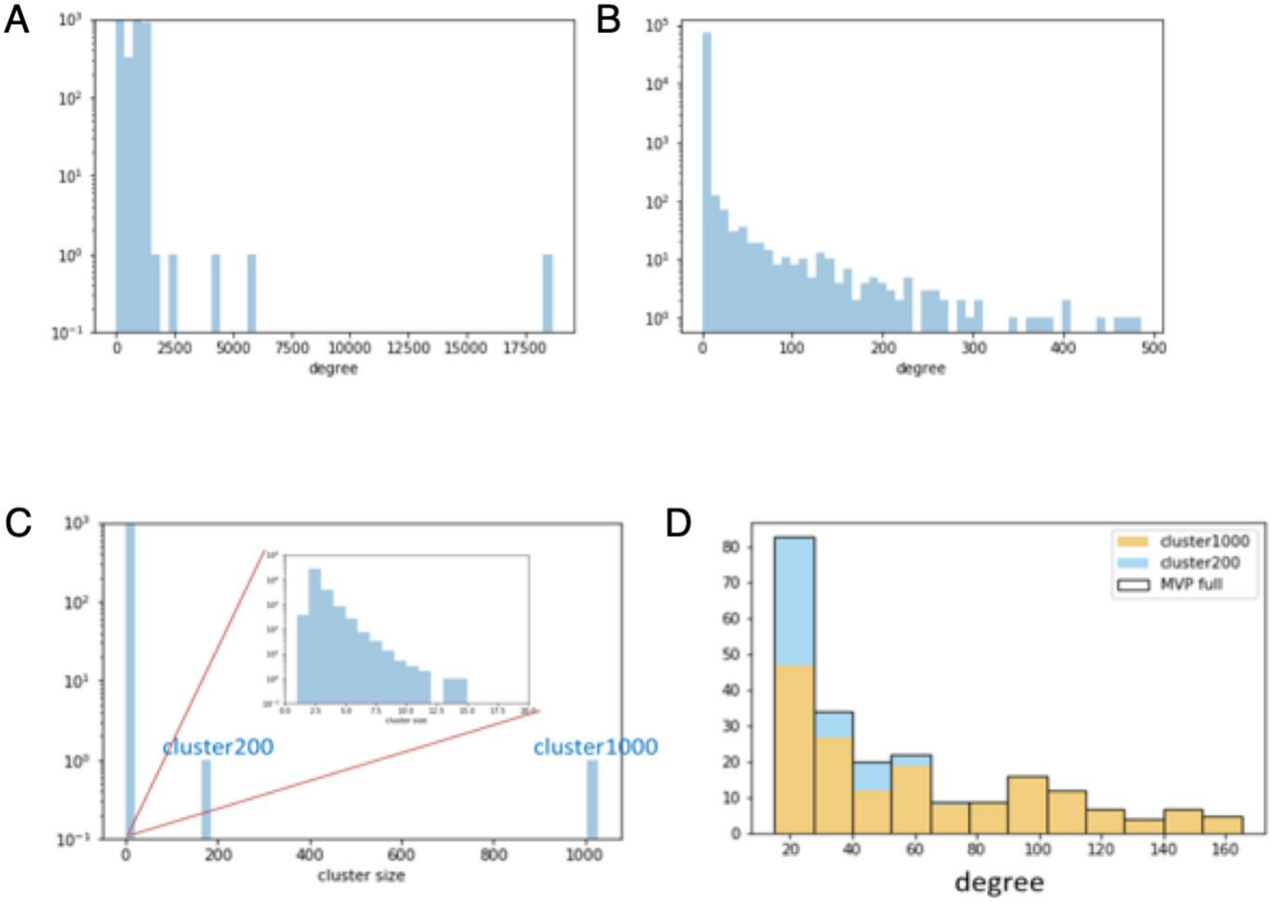


**Figure S10. Analysis of relatedness in the MVP cohort using KING.** (A-B) Number of related individuals per person in (A) round 1 of KING and (B) round 2. (C) Distribution of cluster size of 3rd degree-related individuals after round 2 of KING analysis. The two large clusters of related individuals are denoted as cluster200 and cluster1000. (D) Cluster membership of all individuals with more than 15 3rd degree relatives.

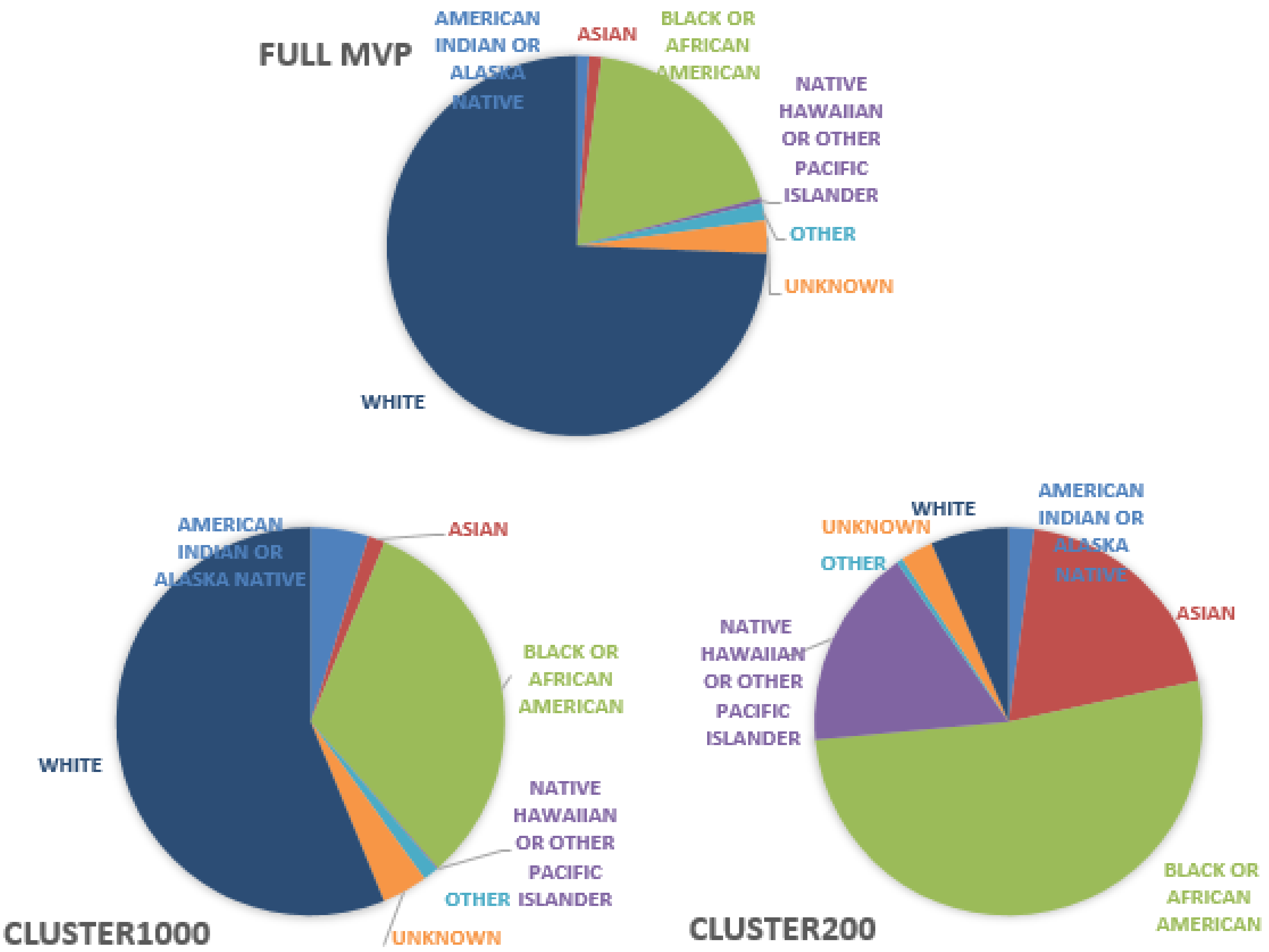


**Figure S11. Race breakdown in the full MVP cohort compared to breakdown in Cluster200 and Cluster1000.** Venn diagrams show proportion of self-identified race in each group.

**Table S1. Breakdown of MVP 1.0 Array markers by chromosome region.**

| Region | Number of markers |
|---|---|
| Mitochondrial chromosome | 270 |
| Non-pseudoautosomal Y chromosome | 142 |
| Pseudoautosomal X and Y chromosome | 1,139 |
| Non-pseudoautosomal X chromosome | 18,026 |
| Autosomal | 667,105 |

**Table S2. Comparison of PLINK-predicted sample gender to EHR-recorded sample gender.**

| Recorded gender / Result | Unknown | Male | Female | OK | Mismatch |
|---|---|---|---|---|---|
| **Unknown** | 10 | 1,646 | 344 | 0 | 2,000 |
| **Male** | 292 | 440,653 | 753 | 440,653 | 1,045 |
| **Female** | 608 | 420 | 41,130 | 41,130 | 1,028 |
| **Total** | 910 | 442,719 | 42,227 | 481,783 | 4,073 |

**Table S3. The top ten ACMG recommended genes for reporting incidental findings, and the number of overlapping MVP 1.0 Array markers.**

| Disease | Gene | Number of markers on the MVP 1.0 array[a] |
|---|---|---|
| Breast-ovarian cancer: familial 2 (MIM 612555) | *BRCA2* | 662 |
| Breast-ovarian cancer: familial 1 (MIM 604370) | *BRCA1* | 541 |
| Marfan's syndrome (MIM 154700) | *FBN1* | 176 |
| Dilated cardiomyopathy 1A (MIM 115200) | *MYBPC3* | 120 |
| Familial hypertrophic cardiomyopathy 4 (MIM 115197) | *MYBPC3* | 120 |
| Familial hypertrophic cardiomyopathy 1 (MIM 192600) | *MYH7* | 101 |
| Brugada syndrome 1 (MIM 601144) | *SCN5A* | 77 |
| Long QT syndrome 3 (MIM 603830) | *SCN5A* | 77 |
| Juvenile polyposis syndrome (MIM 174900) | *SMAD4* | 66 |
| Dilated cardiomyopathy 1A (MIM 115200) | *LMNA* | 60 |

[a] We only included markers located in coding or regulatory regions having ClinVar annotation as either pathogenic or likely pathogenic.

**Table S4. The top 10 disease categories by number of markers shared between the MVP 1.0 Array and the gene-disease associations reported in DisGeNET.**

| Disease | Number of markers | Example diseases |
|---|---|---|
| Congenital, Hereditary, and Neonatal Diseases and Abnormalities | 6,048 | Charcot-Marie-Tooth Disease, Hemophilia A, Hereditary Breast and Ovarian Cancer Syndrome |
| Nutritional and Metabolic Diseases | 4,223 | Diabetes Mellitus, Hyperlipidemia, Obesity |
| Nervous System Diseases | 3,994 | Alzheimer's Disease, Common Migraine, Multiple Sclerosis |
| Cardiovascular Diseases | 3,211 | Abdominal Aortic Aneurysm, Coronary Heart Disease, Hypertensive Disease |
| Neoplasms | 3,087 | Adenocarcinoma, Leukemia, Osteosarcoma |
| Skin and Connective Tissue Diseases | 2,566 | Psoriasis, Rheumatoid Arthritis, Lupus Erythematosus |
| Immune System Diseases | 2,260 | Asthma, Hay Fever, Rheumatoid Arthritis |
| Pathological Conditions, Signs and Symptoms | 2,251 | Chronic pain, motion sickness, seizures |
| Digestive System Diseases | 1,999 | Crohn’s Disease, Inflammatory Bowel Disease, Esophagitis |
| Endocrine System Diseases | 1,985 | Hypothyroidism, Adrenal Cortical Adenoma, Dwarfism |

**Table S5. The Distribution of MVP 1.0 Markers by Minor Allele Frequency, Functional Annotation, and Ancestral Group as Extracted from the ExAC Database.**

| ExAC MAF | | Frameshift | Stop gain/loss | Splice site | Missense | Ins/Del |
|---|---|---|---|---|---|---|
| NFE | MAF ≥ 5% | 28 | 128 | 683 | 15,018 | 26 |
| | 5% > MAF ≥ 1% | 24 | 110 | 246 | 9,288 | 8 |
| | 1% > MAF > 0% | 1,533 | 10,805 | 4,260 | 138,093 | 94 |
| | MAF = 0% | 2,344 | 8,314 | 2,554 | 39,190 | 77 |
| AFR | MAF ≥ 5% | 522 | 2,670 | 1,296 | 33,791 | 40 |
| | 5% > MAF ≥ 1% | 37 | 230 | 522 | 19,664 | 21 |
| | 1% > MAF > 0% | 718 | 3,442 | 2,750 | 87,686 | 64 |
| | MAF = 0% | 2,619 | 13,014 | 3,175 | 60,445 | 80 |
| EAS | MAF ≥ 5% | 27 | 118 | 592 | 14,058 | 22 |
| | 5% > MAF ≥ 1% | 24 | 79 | 174 | 6,453 | 5 |
| | 1% > MAF > 0% | 915 | 4,304 | 1,662 | 35,304 | 47 |
| | MAF = 0% | 2,929 | 14,855 | 5,315 | 145,766 | 131 |
| SAS | MAF ≥ 5% | 32 | 127 | 650 | 14,934 | 23 |
| | 5% > MAF ≥ 1% | 13 | 92 | 232 | 7,279 | 9 |
| | 1% > MAF > 0% | 605 | 4,394 | 2,272 | 69,268 | 38 |
| | MAF = 0% | 3,243 | 14,744 | 4,589 | 110,103 | 135 |
| AMR | MAF ≥ 5% | 29 | 131 | 633 | 14,932 | 23 |
| | 5% > MAF ≥ 1% | 22 | 74 | 263 | 8,090 | 11 |
| | 1% > MAF > 0% | 401 | 2,907 | 2,490 | 101,308 | 84 |
| | MAF = 0% | 3,443 | 16,244 | 4,357 | 77,243 | 87 |
| Not in ExAC | | 4,513 | 5,091 | 680 | 10,985 | 168 |

**Table S6. MVP 1.0 Array markers categorized by PolyPhen and SIFT score categories based on predicted protein sequence changes.**

| | | SIFT[a] | | | |
|---|---|---|---|---|---|
| | | **deleterious** | **deleterious Low confidence** | **tolerated** | **tolerated low confidence** |
| **PolyPhen[a]** | **probably_damaging** | 47,954 | 3,005 | 12,447 | 716 |
| | **possibly_damaging** | 14,107 | 1,778 | 14,576 | 1,160 |
| | **benign** | 13,355 | 2,043 | 74,197 | 7,950 |
| | **unknown** | 752 | 612 | 1221 | 798 |

[a]PolyPhen and SIFT scores and categories were obtained from the Ensemble Variant Effect Predictor.

**Table S7. MVP 1.0 array imputation accuracy for the five 1000 Genomes Project Phase 3 continental populations.**

| | 1% <MAF <5% | | MAF ≥ 5% | |
|---|---|---|---|---|
| **Population** | **Number of imputed markers** | **Accuracy (mean.$r^2$)** | **Number of imputed markers** | **Accuracy (mean.$r^2$)** |
| EUR | 2,902,508 | 0.777 | 7,244,340 | 0.931 |
| AFR | 7,339,754 | 0.767 | 10,097,459 | 0.874 |
| EAS | 2,297,176 | 0.656 | 6,693,655 | 0.890 |
| SAS | 3,190,047 | 0.710 | 7,437,310 | 0.904 |
| AMR | 4,140,903 | 0.824 | 7,464,928 | 0.925 |

**Table S8. Breakdown and number of closely related pairs in the MVP Cohort and UK Biobank.**

| | **MVP** | **UK Biobank** |
|---|---|---|
| Total sample size | 459,777 | 488,410 |
| Total pairs (billions) | 105.70 | 119.27 |
| Trios | 28 | 1,066 |
| Monozygotic twins | 49 | 179 |
| Parent-child | 2,886 | 6,276 |
| Full sibling | 5,330 | 22,666 |
| 2nd or 3rd degree | 9,055 | 78,041 |

### Supplemental Materials and Methods

#### Axiom Array Terminology

Here, we first define terms used in standard marker quality control. A marker refers to a genetic variation at a specific genomic location in the DNA of a sample. Both single nucleotide polymorphisms (SNPs) and insertion/deletions (indels) are markers and are genotyped. A set of one or more probe sequences from which array intensities are combined to interrogate a marker is referred to as a probeset. Most Axiom® markers are interrogated with one or two probesets derived from the forward strand sequence, the reverse strand sequence, or both.

#### Thermo Fisher Scientific Axiom® Genotyping Platform

The MVP 1.0 array is based on the Applied Biosystems Axiom® Biobank core variants. The Axiom® Biobank array modules (**A-F1**) include:

(1) Genome-wide coverage for common European variants *(N= 246,038; Module "High coverage imputation grid")*: Markers were selected using Affymetrix imputation aware marker choice algorithms to provide genome-wide coverage in European populations of common (MAF ≥ 5%) markers using the CEU panel from 1000 Genomes Project.
(2) Loss-of-function (LoF) SNPs and indels *(N= 70,396; Module "High confidence LoF SNPs and Indels")*: Markers with a high-confidence loss of function call, possible splice-site impact, or representing known, disease-associated mutations. Markers are extracted from an exome sequencing initiative of 26,000 people. This category of markers also includes known disease-causing mutations and potential splice variants from Human Gene Mutation Database (HGMD).
(3) Exome SNPs and indels *(N= 264,193; Module "Exome Grid")*: Non-synonymous coding markers in exonic regions of the genome. This content was derived from the Exome Chip Design Consortium, the NHLBI Exome Project, the Genetics of Type 2 Diabetes program (GoT2D), the 1000 Genomes Project, the Cancer Genome Atlas Project, the SardiNIA Medical Sequencing Study, the Autism Exome Sequencing Study, the UK10K project, and others.
(4) Expression quantitative trait locus *(N= 5,884; Module "Expression quantitative loci from GTEx")*: Markers with known associations to RNA expression traits taken from NCBI Genotype-Tissue Expression (GTEx) database.
(5) Pharmacogenomic/ADME *(N= 2,031; Module "Pharmacogenetics")*: Markers associated with phases of drug absorption, distribution, metabolism, and excretion (ADME). This content was derived from the PharmaADME and PharmGKB databases. The content is enriched for markers in Clinical Pharmacogenetics Implementation Consortium (CPIC) guidelines and those used in calling "star alleles" variants of genes that affect drug metabolism.
(6) *ApoE (N=2, not shown)* The two SNPs, rs429358 and rs7412, which define the ApoE isoforms known to be associated with risk of Alzheimer's disease and lipoprotein-associated conditions.

##### MVP-Specific Modules

(1) SNPs and indels based on known associations with diseases or traits of interest (N=21,627; module "*Other diseases associated and exome markers*"). Over 800 potential clinical conditions and 3,400 gene regions are covered by this module with an emphasis on

conditions of high prevalence in US Veterans such as substance abuse[1], amyotrophic lateral sclerosis (ALS [MIM: 105400]) [2], chronic obstructive pulmonary disease (COPD [MIM: 606963])[3], and diabetes[4]. Additionally, we included the 59 genes currently recommended for reporting of incidental findings by the American College of Medical Genetics and Genomics (ACMG)[5,6] in anticipation of future clinical applications.

(2) Human leukocyte antigen (HLA) and killer-cell immunoglobulin-like receptor (KIR) region markers for fine mapping and imputation (N=8,058; module *"Markers within genomic regions of interest")*. Genes in the *HLA* region are critical players in immune function, and some have known roles in histocompatibility[7,8]. We also included additional *KIR* markers that have an important role in infectious disease, autoimmune disorders, cancer, and reproduction by virtue of the region's interaction with the *HLA* region. Furthermore, to optimize coverage and improve imputation quality in non-European ancestry populations, we also included additional *HLA* and *KIR* markers in multiple ethnicities, including African ancestry (N= 1,043; Module "HLA imputation booster for non-Caucasian cohorts").

(3) Psychiatric condition-associated markers (N=26,928, module *"Psychiatric relevant markers"*). 11-20% of US military personnel who served in Operations Iraqi Freedom and Enduring Freedom are estimated to have diagnosed or undiagnosed post-traumatic stress disorder (PTSD)[9]. The lifetime prevalence of PTSD among Vietnam Veterans, who form the largest subset of the MVP cohort, is 15-20%[10]. To study genetic factors that affect the development of mental illness and the functional disability and cognitive impairment that accompany these conditions, we included additional markers extracted from the Psychiatric Genomics Consortium[11] (personal communications with PGC group leaders) that have been linked with common psychiatric disorders such as schizophrenia (MIM: 181500), bipolar disorder (MIM: 125480), autism spectrum disorders (MIM: 209850), attention deficit hyperactivity disorder (MIM: 143465), major depressive disorder (MIM: 608516), obsessive compulsive disorder (MIM: 164230), anorexia (MIM: 606788), and Tourette syndrome (MIM: 137580). Many of these variants are rare for some ethnicities and have minor allele frequency (MAF) < 1%.

(4) Candidate genes associated with rheumatoid arthriti*s* (MIM: 180300) (N=11,144; module *"Rheumatoid Arthritis relevant markers"*). Markers include a) rheumatoid arthritis-associated SNPs outside of the MHC region[12,13,14] (personal communications); b) SNPs with minor allele count (MAC) > 2 found in genes in rheumatoid arthritis risk loci in the National Heart, Lung, and Blood Institute (NHLBI) Exome Sequencing Project (ESP) data (European ancestry cohort); and c) SNPs in the 1000 Genomes Project European population that overlap with the rheumatoid arthritis risk loci having MAF > 1% after linkage disequilibrium pruning to remove SNPs with $r^2$ >0.8.

(5) Genome-wide imputation booster for African ancestry population (N=50,000; module *"Imputation booster"*). The MVP population contains multi-ethnic groups, and African Americans comprise the second largest self-reported ethnic group in the MVP. To optimize the coverage of African genetic ancestry, an additional set of markers were tiled on the MVP array to boost corresponding imputation coverage. Markers were selected using Axiom® imputation aware marker choice algorithms[15] and the YRI panel from the 1000 Genomes Project to provide genome-wide coverage in African populations of common (MAF ≥ 5%) markers.

#### Overlap with databases of functional and disease-related variants

The MVP 1.0 array was designed to increase coverage of known functional and disease- or trait-associated variants across the genomes of diverse populations. To evaluate this coverage, we aimed to check the coverage of MVP 1.0 markers with markers from various data sources cataloging potential functional variants in multiple populations that were not considered during the design of the chip: the NHGRI-EBI GWAS catalog[16,17], ClinVar [18]; the Exome Aggregation Consortium (ExAC)[19,20]; and DisGeNET, which also contains the genes in the ACMG guideline recommendations for reporting incidental findings and gene-disease associations[5,6]. Markers in the MVP 1.0 chip overlapped with 7,199 variants in the GWAS catalog v1.0[16], 31,067 in ClinVar[18], and 294,092 in ExAC r.0.3.1[20]. As expected by the number of GWAS hits in the GWAS catalog, the top five diseases associated with variants shared between MVP 1.0 and the GWAS catalog are schizophrenia, rheumatoid arthritis, Crohn disease, type 2 diabetes, and systemic lupus erythematosus, and the top five biological traits are height, BMI, HDL, cholesterol, LDL cholesterol, and triglycerides. Furthermore, around half of the variants shared between MVP 1.0 and ClinVar (17,124 of 31,067) are classified by ClinVar as pathogenic or likely pathogenic and have low allele frequency in all ancestral groups.

Additionally, MVP 1.0 contains markers overlapping with pathogenic variants residing in all 59 ACMG genes[5,6]. The top ten ACMG genes and the corresponding number of MVP 1.0 markers are listed in Table S3. DisGeNET also contains one of the largest publicly available collections of genes and variants associated with human diseases[21], and the MVP 1.0 array contains 6,048 DisGeNET markers associated with congenital, hereditary, and neonatal diseases and abnormalities, which represents the largest overlapping disease category. The top ten DisGeNET categories and the corresponding number of MVP 1.0 markers are listed in Table S4.

Variants in the ExAC database predicted to have severe consequences (e.g. missense, frameshift, stop-gain, start-loss variants) were highly enriched in the rare and low frequency MAF ranges (Table S5). Furthermore, in addition to containing markers for variants observed in the 1000 Genomes Project and ExAC database, the Axiom Biobank Genotyping Array backbone has many variants not found in these databases. For example, more than half of the frame shift variants and one-third of the insertion or deletion variants were absent from these databases (Figure S1). We also examined MVP 1.0 nonsynonymous single nucleotide variants (nsSNVs) in detail by implementing functional annotation using the Ensembl Variant Effect Predictor (VEP), release 86[22]. 66,790 coding variants were predicted as deleterious by both PolyPhen[23] and Sift[24] (Table S6), and 16,598 non-coding variants were annotated as regulatory variants.

Finally, we examined the allele frequency distribution of variants on the MVP 1.0 chip to assess its coverage across ancestral population groups. Allele frequencies of variants in the GWAS catalog and ClinVar were assessed using 1000 Genomes, whereas ExAC variant frequencies were used directly. Four MAF categories were defined as: common variants (MAF ≥ 5%), low-frequency variants (MAF 1-5%), rare variants (MAF < 1%), and monomorphic variants (MAF = 0%). In general, markers in the MVP 1.0 chip showed similar distribution of MAFs across all ancestral groups in all subsets (Figure S2). As expected, most markers in the MVP 1.0 chip that overlaps NHGRI-EBI GWAS catalog variants were common (MAF ≥ 5%, Figure S2A**)**, whereas markers that overlap ClinVar variants were mostly uncommon (MAF < 5%, Figure S2B**)**. On the other hand, although the proportion of common variants in ExAC was fairly consistent across ancestral groups, the proportion of less common ones showed greater variability (Figure S2C).

#### Assessing imputation accuracy of the MVP v1 chip

Imputation was executed using IMPUTE29 and the 1000 Genomes Phase 3 reference panel10. To assess imputation accuracy, we performed ten-fold cross-validation, which corresponds to a sub-sample of the participants in each of the five populations. The 1000 Genomes Phase 3 genotypes of the held-out samples for the markers on the MVP 1.0 array were used to impute the genotypes of the markers in the modified reference panel. The accuracy of imputed variants was calculated as the squared Pearson correlation coefficient (r2) between imputed genotype dosages (0–2) and the genotypes from the 1000 Genomes Phase 3 release (0,1,2). The results were stratified into two non-overlapping minor allele frequency (MAF) bins MAF ≥ 5% and 1% < MAF < 5%, and accuracy was summarized as the mean r2 over the marker count in the MAF bin.

In addition to the aim of increasing coverage of functional variants, the MVP 1.0 array was designed to maximize imputation performance for GWAS studies. We analyzed imputation performance of the array on 1000 Genomes Project data by performing cross-validation with 10% of the samples at a time. We report the accuracy of imputed variants across two MAF bins in terms of the squared Pearson correlation coefficient (r2) between imputed genotype dosages (0-2) and the genotypes from the 1000 Genomes Project Phase III release (0, 1, 2) (discussed in Materials and Methods.) (Table S7). For example, the mean r2 for the EUR group is 0.931 in common variants and 0.777 in low frequency variants. Among the five ancestry groups, imputation of common variants was most accurate for the EUR population, while imputation of less common variants was most accurate for the AMR population. The high accuracy of imputation of less common variants in the AMR population reflects the admixture from European, African, and Native American ancestries, two of which were already included in the 1000 Genomes Project.

#### Vendor quality control (QC)

Two vendors, referred to as Vendor 1 and Vendor 2, performed initial genotyping of MVP samples for internal QC purposes. The vendors genotyped all MVP samples on the custom MVP 1.0 array. To batch samples in preparation for genotype calling, the vendors grouped approximately 5,000 samples into each batch and used either the Axiom® Analysis Suite (Windows graphical user interface) or Affymetrix® Power Tools (APT, command line software) to execute the *AxiomGT1* genotyping algorithm. Vendor 1 called genotypes with the Axiom® Analysis Suite versions 1.15, 1.16, 2.3, and 2.4. Vendor 2 called genotypes with the APT software and used versions 1.15.1 and 1.16.1. Both the Axiom® Analysis Suite and the APT software call genotypes in each batch independently of other batches.

The vendors followed the Axiom® Best Practices to filter the data for QC purposes[25] and so reprocessed samples with dish QC (DQC) values less than 0.82 or sample QC call rate less than 97%. Whereas Vendor 2 used a threshold of 97% to filter samples in all batches for re-genotyping, Vendor 1 began with a threshold of 97% but switched to a 98% threshold after genotyping several batches to increase genotyping quality. This is the reason why Vendor 1's QC call rate is consistently higher than that of Vendor 2 even after re-batching and re-genotyping on the full cohort (Figure S5 A-C). Only samples that pass internal vendor QC were sent to the VA.

The vendors also investigated plates with the plate QC metric (average QC call rate of passing samples over the plate) lower than 98.5% to identify any systematic row or column failure and removed these plates if found. Since both vendors re-processed failed samples immediately after performing the

Best Practices analyses and did not always return the failed samples in the delivered data, we relied on the vendors' reports for the plate QC, sample DQC, and QC call rate values.

#### Genotype calling on the MVP cohort

All samples that passed vendor QC were then re-batched and recalled internally at the VA under a unified workflow. After comparing vendor and multiple additional batching strategies, we separated MVP samples by vendor and then batched samples in groups of 4,000 to 5,000 samples by each sample's plate scan date. We incorporated samples into each batch at the unit of a plate so that all samples on one plate would be within the same batch. Since experimental metadata captured in *CEL* files (Axiom® files that report the array intensity data) is considered the most reliable, we extracted the plate scan date from the CEL file header by using an APT program called calvin-extract. After batching samples, we had 112 batches with a median of 4,500 samples. Then, we called genotypes in each batch with the APT software, version 1.18, following the Axiom® Best Practices Genotyping workflow[25].

#### Initial QC

To provide an initial assessment of MVP sample genotyping quality, we analyzed several QC metrics in detail:

1) The DQC metric is an approximate measure of contrast between genotyping channels at non-polymorphic genomic locations[25]. Because the DQC metric is calculated only on non-polymorphic probesets, this metric assesses the signal to background noise ratio for each channel and should be close to 1 for high-quality samples.
2) We also analyzed QC call rates (call rates computed over a set of 20K QC probesets, also referred to as step 1 call rate) and sample call rates over all probesets (also referred to as step 2 call rate) by batch for all samples. The sample call rate is similar except it is calculated on genotype calls over all probesets in a second round of calling after samples that failed the QC call rate threshold were removed.
3) The final metric, sample missingness, is the fraction of genotype calls which are missing per individual after the advanced marker QC procedure over the number of markers that are called in at least one individual in the sample. The sample missingness is an indicator of sample genotype call completeness after the conclusion of all QC procedures.

#### Standard marker quality control

Standard marker quality control is executed by the Axiom® Best Practices Genotyping workflow. This workflow executes rules to select a *best* probeset for markers that are interrogated with more than one probeset, which is followed by the sorting of probesets into four *recommended* and three *not-recommended* SNP QC classes. Three recommended classes require probesets to produce well-resolved genotype clusters for probesets with three (the *PolyHighResolution* class), two (the *No Minor Homozygous* class), or one (the *Mono High Resolution* class) genotype clusters in the given batch (see the Axiom® Genotyping Solution Data Analysis Guide). The fourth recommended class is "Hemizygous" for Y chromosome and mitochondrial markers. The best practices guideline is to only use genotype calls from a given batch for marker probesets that are both *best* and *recommended*.

#### Advanced Marker Quality Control Procedure for the MVP 1.0 array

##### Approach 1: Exclude probesets that fail Advanced QC tests

The goal of the Advanced QC tests is to identify probesets that systematically underperform in most batches or that are interrogating multi-allelic markers. Such probesets were identified in five ways: (1) probesets for multi-allelic markers; (2) probesets with higher than expected variance in allele frequency between batches; (3) probesets that failed to cluster into recommended classes in a large fraction of batches; (4) probesets with large differences in allele frequency when compared to reference datasets; and (5) mitochondrial or Y chromosome probesets that fail visual inspection of their cluster plots.

###### *Approach 1.1: Probesets for multi-allelic markers*

Because the current *AxiomGT1* genotyping algorithm only calls the three genotypes for bi-allelic markers, probesets for multi-allelic markers, which have a minimum of six genotypes, are excluded. Only 0.22% of probesets interrogate multi-allelic markers.

###### *Approach 1.2: Probesets with inconsistent allele frequencies across batches*

Probesets exhibiting larger than expected variance in allele frequency across genotyping batches could indicate genotype calling issues. We used a statistic called the ratio A allele frequency (RatioAAF) to measure the stability of a probeset's allele frequency across batches[26]. The RatioAAF is defined as the expected allele frequency variance over the observed allele frequency variance:

$$RatioAAF = \frac{p'(1-p')}{\mathrm{var}(P)}$$

where $P$ is a vector consisting of the marker allele frequencies across all recommended batches and $p'$ is the average allele frequency across all recommended batches. A larger RatioAAF indicates a more stable allele frequency. Overall, 3.41% probesets have RatioAAF less than 1,000 when computed over recommended batches. These probesets are often recommended in individual batches but produce inconsistent allele frequencies when measured over multiple batches. Visual inspection of a sampling of cluster plots of low RatioAAF probesets confirms that such probesets produce poor genotype clusters.

###### *Approach 1.3: Probesets that are recommended in few batches*

Probesets that are not often classified into one of the four recommended categories (discussed above) over most batches are typically genotyping poorly. Visual inspection of cluster plots for probesets recommended in less than 20 batches out of 112 confirms that such probesets produce poor genotype clusters and suggests that probesets recommended in only a few batches are fundamentally not working. 2.68% of probesets were eliminated by this filter.

###### *Approach 1.4: Probesets with allele frequencies diverging from those in reference datasets*

A small fraction of probesets demonstrate large discrepancies in reference allele frequency between the MVP dataset and the 1000 Genomes Project, phase 1 CEU reference population. Such probesets were identified with the following thresholds on the reference allele frequencies: (MVP $< 0.02$ AND CEU $\geq 0.5$) OR (MVP $> 0.98$ AND CEU $\leq 0.5$). Allele frequency thresholds were set more stringently for the MVP samples because of the larger sample size. Only 0.82% of probesets were excluded by these criteria.

***Approach 1.5: Probesets on the mitochondrial or Y chromosome that fail visual inspection***

We conducted a manual review of cluster plots for probesets on the mitochondrial and Y chromosomes as recommended by the Best Practices for these hemizygous markers. Only a small fraction of probesets (0.01%) were excluded by this filter.

**Approach 2: Exclude calls from a mis-clustering event by batch**

After removing probesets that were fundamentally not working over all genotyping batches, we then turned to a batch level metric of performance. If a probeset was generally working but not one of the four recommended classes in a particular batch, the calls for that probeset in that batch were set to missing. In this way, we retained working probesets while removing calls in sporadic problematic batches.

**Approach 3: Select one probeset per marker over all batches**

There are 36,601 markers that are interrogated by more than one probeset on the MVP array. For markers that are interrogated by more than one probeset, the Axiom® Best Practices Workflow designates only one probeset as the best probeset per batch based on twelve performance metrics calculated as a part of the genotype calling procedure. However, the probeset that is designated as the best probeset can be different in different batches. We used five metrics to determine the best probeset per marker over all batches. These five metrics included the number of batches in which the probeset succeeded, the number of batches in which the probeset was marked best, the number of batches in which the probeset was classified as Poly High Resolution, the average probeset call rate over all batches, and the average Fisher Linear Discriminate (FLD) over all batches. We compared each probeset's value for each of these metrics sequentially and selected the first probeset that had a higher value for one of the metrics. If all probesets for a marker were equivalent for all five metrics, we selected the first probeset in alphanumeric order.

#### Correction of batch variation

Individuals in subsequent tranches (Release A and Release B) were randomly sampled from the same population and similarly processed. Therefore, we hypothesized no common genetic variants were significantly associated with assignment to release. We tested this hypothesis by conducting a preliminary GWAS. We restricted the GWAS to samples with European ancestry (GBR >= 0.9 in ADMIXTURE). We used genotypes called from both tranches combined and assigned samples to pseudo-control and case groups according to their Release tranches. Specifically, we assigned Release A samples to the control group (n=238,229) and Release B samples to the case group (n=73,153) as shown in Figure S4B. We ran the association using only common markers (MAF > .05) and used the first 10 principal components (projected to 1000 Genomes Project principal components) as covariates and a linear-mixed model association in BOLT-LMM, which adjusts for cryptic relatedness.

We found 204 significant ($p \leq 5 \times 10^{-8}$) genome-wide independent signals (Figure S4D). The signals were single variant associations without local LD structure based on manual inspection of locuszoom plots[27], suggesting batch effects. We subsequently ran an association adjusting for batch membership as a categorical variable (n = 112) and found only one genome-wide significant hit (in the HLA region). We also ran an association randomly assigning each individual to case or control and observed no significant associations.

For each variant that was genotyped, we then used Fisher's Exact Test to identify batches in which allele distributions significantly differed from that of all other batches. This tests the null hypothesis that a given batch has the same genotype frequencies as all other batches combined. We compared distribution of allele frequencies across batch-SNP combinations (~60 million) with significant deviance defined as $p < 6.7 \times 10^{-10}$ (i.e., 0.05/batch-SNP combinations). 173 of the 204 genome-wide significant SNPs from the Bolt-LMM GWAS failed the Fisher's Exact Test. An example of a SNP-batch combination that was detected by the Fisher's Exact Test is shown in Figure S4A. We then set all failing SNP-batch combinations to 'no-call' (approximately ~50,000 of ~60,000,000 batch-SNP combinations) and re-ran the Bolt-LMM GWAS as above, this time finding 73 significant independent signals.

We then investigated the remaining 73 significant hits by examination of genotype cluster plots. We identified certain patterns for which Fisher's Test would fail and came up with a series of additional batch correction rules. For probes that fail Fisher's Test in at least one batch:

1) If the RatioAAF is less than 1,000, then exclude all batches for the probeset.
2) If the batches with passing probesets all have fewer clusters than the batches with failing probesets, then switch the pass/fail assignment.
3) If the batch has a call rate less than 98% and FLD less than 5, then fail the batch.
4) If a batch is classified as No Minor Homozygous and has an HWE p-value $< 10 \times 10^{-4}$, then fail it.

An overview of these steps and number of SNPs identified is summarized in Figure S4C**.**

After applying these additional batch correction rules, we re-ran the Bolt-LMM GWAS and found an overall reduction in the number of significant associations from 204 to 47 (Figure S4 D-E). These remaining associations were unlikely to be meaningful signals based on manual visual inspection of locuszoom plots and were set to no-call.

#### Relatedness analysis

Even after excluding 35 individuals with more than 200 relatives, we observed two clusters (denoted as cluster200 and cluster1000) with more than 100 family members in our KING analysis (Figure S10 C-D). Interestingly, all individuals with greater than 15 3$^{rd}$ degree relatives are in one of the two clusters. Members of each cluster are tightly interconnected with each other rather than there being a few individuals promiscuously interacting with other members of the cluster. It is unlikely that these two clusters are real families, and further investigations are warranted. We also observe an underrepresentation of individuals self-identifying as "white" in these two clusters (Figure S11), suggesting the low weight SNP strategy may not have fully removed the overestimation of relatedness in admixed individuals. We flagged all individuals in cluster200 and cluster1000 and redefined a new set of unrelated individuals using the largest_independent_vertext_sets() function to identify the largest set of unrelated individuals to use in our analyses.

#### Web Resources

Axiom Genotyping Solution Data Analysis Guide, https://assets.thermofisher.com/TFS-Assets/LSG/manuals/axiom_genotyping_solution_analysis_guide.pdf

Locuszoom, http://locuszoom.org/
